# The avocado pangenome reveals dynamic clustering and lineage-specific diversity of *NLR* genes

**DOI:** 10.1101/2025.10.28.684993

**Authors:** Robert Backer, Alicia Clarke, Alicia Vermeulen, Aureliano Bombarely, Noёlani van den Berg

**Affiliations:** Department of Biochemistry, Genetics and Microbiology, University of Pretoria, Pretoria, Gauteng, South Africa; Hans Merensky Chair in Avocado Research, Forestry and Agricultural Biotechnology Institute, University of Pretoria, Pretoria, Gauteng, South Africa; Instituto de Biología Molecular y Celular de Plantas, Consejo Superior de Investigaciones Científicas- Universitat Politècnica de València (IBMCP-CSIC-UPV), Valencia, Spain

## Abstract

Avocado is an economically important perennial tree crop with a complex domestication history, yet modern genomic resources are limited. We present the first high-quality avocado pangenome, spanning seven diverse accessions across Mexican, Guatemalan, and West-Indian lineages. PacBio HiFi, partially phased assemblies deliver near chromosome-level continuity (N50 ~57 Mb; BUSCO >96%), and functional annotations show enrichment of immune-related families. Core–accessory partitioning indicates adaptive functions — pathogen response and secondary metabolism — are overrepresented in accessory genes. Nucleotide-binding Leucine-rich repeat (*NLR*) genes were catalogued (226–256 per accession), showing lineage-specific expansions, diverse domain architectures, and frequent chromosomal clustering. Structural variants concentrate within *NLR* loci, marking hotspots of pathogen-detection diversification. Comparative analyses show ~54% of *NLRs* are shared, with extensive functional and sequence diversity among accessions. Together, these results define the avocado NLRome and its core– accessory interplay, providing a graph-based framework to accelerate discovery of resistance loci and breeding for durable disease resistance.

## Introduction

Genomic resources for perennial tree crops trail those of annuals, hampered by biological and technical challenges. Many fruit and nut trees are long-lived, with extended juvenility and long generation times, making conventional breeding arduous^1–4^. Their genomes are often large, structurally complex, and highly heterozygous from outcrossing, complicating *de novo* assembly^5^. Consequently, whereas annuals such as rice^6^, common bean^7^, and maize^8^ have large pangenomes, most tree crops lack comparable references. Notably, perennials such as apple^9^, peach^10^ and citrus^11^ only recently moved beyond single references to first pangenomes. This gap hampers discovery of structural-variants (SVs) and trait-linked alleles in tree crops^12–14^, underscoring the need for deeper catalogues of perennial diversity.

Avocado (*Persea americana*) exemplifies these challenges. Cultivated avocados derive from three landraces – Mexican, Guatemalan, and West Indian – but genomics focused on a few elite cultivars within the admixed Hass lineage^15–17^. The first reference genome was only published in 2019^16^, followed with a chromosome-scale Hass assembly in 2022^17^. However, no pangenome captures diversity across landraces and hybrids. This need is evident from studies of defence-related genes^18–20^, reporting large gene-number differences across assemblies, highlighting the need for an avocado pangenome.

Variation in defence-gene sequences can influence immune activation and disease outcome^21–23^. Many defence proteins activate immunity upon recognising pathogen-associated patterns or pathogen effectors. Effector recognition is highly specific; thus, sequence differences may alter protein function^24^. Nucleotide-binding Leucine-rich repeat (NLR) receptor proteins, for example, mainly recognise effectors via a C-terminal Leucine-rich repeat region^25^. Mutations within this domain have been linked with pathogen susceptibility^26^. Furthermore, *NLR* copy number variants are frequently associated with quantitative trait loci linked to disease resistance^27–29^. Understanding population-level *NLR* variation will clarify plant-pathogen interactions, resistance, and the evolutionary pressures shaping this protein family^30^.

*NLRs* often occur in clusters — homogeneous tandem arrays or heterogeneous groups from segmental duplication – that serve as evolutionary hotspots, generating immune diversity^30^. Pangenome approaches are therefore valuable, capturing conserved and variable *NLRs* across avocado accessions and enabling comprehensive study repertoire and lineage-specific diversification^12–14^.

Here we present the first high-quality, partially phased avocado pangenome spanning all major horticultural lineages — a foundation for trait discovery and improvement. We delineate core and accessory gene sets; the accessory compartment is enriched for defence- and secondary-metabolism GO terms and shows expansions of NLR-linked Pfam domains. Using a graph pangenome, we map genome-wide structural variation, with inversions, deletions and insertions enriched in *NLR*-associated regions. Comparative analyses reveal differences among accessions in NLR domain organisation and sequence conservation, providing candidates for functional validation. The identification of accession-specific variants also offers candidate NLRs for functional validation. Ultimately, this pangenome reveals extensive NLRome diversity and lays the groundwork for durable resistance strategies.

## Results

### Nuclear assemblies

We generated high-quality, partially phased genome assemblies for seven *P. americana* accessions, including the previously published Hass genome^17^, spanning the major landraces and hybrid groups. PacBio HiFi sequencing on Revio yielded ~26–36 Gb per accession (mean read length >15 kb, Q>27). Assemblies built with Hifiasm produced primary and alternate haplotypes; primaries were the most contiguous. After filtering organelle and other non-nuclear sequences, primary assemblies ranged from 862 Mb (Leola™) to 915 Mb (Hass; Table S1). Contig counts varied from 56 (Choquette) to 810 (Ashdot; Fig. S1), yet L90 values of 16 to 28, indicated that most of each assembly was captured in relatively few contigs. N50 values ranged from 3.6 Mb (Hass) to 57 Mb (Ashdot).

For the previously published Hass dataset, available raw data (~17.5 Gb) were substantially lower than the ~47.9 Gb reported in the original study^17^, likely contributing to its reduced contiguity relative to the newly sequenced accessions. BUSCO completeness for Hass (96.7%) matched our assemblies (96.3–96.7%). Merqury confirmed comparable quality, with Hass QV 61.3 and completeness of 98.0% versus QV 63.3–71.6 and completeness of 98.8–99.3% for the newly sequenced accessions (Table S1).

To assess structural consistency, primary assemblies were aligned to the West-Indian (WI) pure accession genome. Dotplots showed broad syntenic concordance with limited lineage-specific rearrangements (Fig. S2). The main exception was Choquette contig 0003, in which WI chromosome 6 had been appended to chromosome 4; we corrected this misjoin by manually splitting the contig. Aside from this case, structural differences were largely confined to regions of lower concordance — most notably central chromosomal regions — none of which could be confidently classified as assembly errors.

Very few contigs showed concordance with chromosome 0 of the WI reference (Fig. S2). Earlier assemblies used “chromosome 0” for unplaced sequences that often represented contaminants, organelle fragments, or other artefacts. Our workflow used stringent contaminant and organelle filtering and, without explicitly targeting “chromosome 0”, effectively excluded such sequences.

### Organelle assemblies

Complete circular chloroplast genomes were recovered for all seven accessions (152,609– 152,763 bp; Ashdot; Fig. S3, S4, Table S2), consistent with prior reorts^31^. Mitochondrial assemblies were more variable; Hass, Ashdot, and Mike each yielded two circular contigs totalling ~850 kb, similar to the recently published complete *P. americana* mitogenome^32^. Dusa^®^, Choquette, and Leola™ produced near-complete but non-circular assemblies, and no contiguous mitogenome was reconstructed for Gottfried. This pattern likely reflects nuclear DNA isolation minimizing organellar carry-over.

### Repeat landscape and transposable element composition

All assemblies displayed similar repeat architectures, with repetitive DNA comprising ~59– 60% of the genome space (Fig. S5; Table S3). Long terminal repeat (LTR) retrotransposons dominated (~35%): ~18–19% LTR/unknown (unclassified), ~10–11% Gypsy, and ~5% Copia. DNA transposons accounted for ~10–11% (primarily Mutator-like, ~5–6%, and hAT, ~2%), long interspersed nuclear elements (LINEs) for ~6–7%, and miniature inverted-repeat transposable elements (MITEs) and other low-complexity repeats contributed <5%. Approximately 3% of repeats could not be assigned to known families.

Divergence (Kimura) landscapes revealed a pronounced wave of recent LTR expansion peaking at ~10% divergence, largely driven by unclassified and Gypsy elements, with broader shoulders indicating older bursts (Fig. 1). Temporal patterns were highly conserved across accessions: overlaid landscapes (Fig. 1h) were nearly indistinguishable, indicating shared historical retrotransposon activity rather than recent accession-specific proliferation.

**Figure 1.**
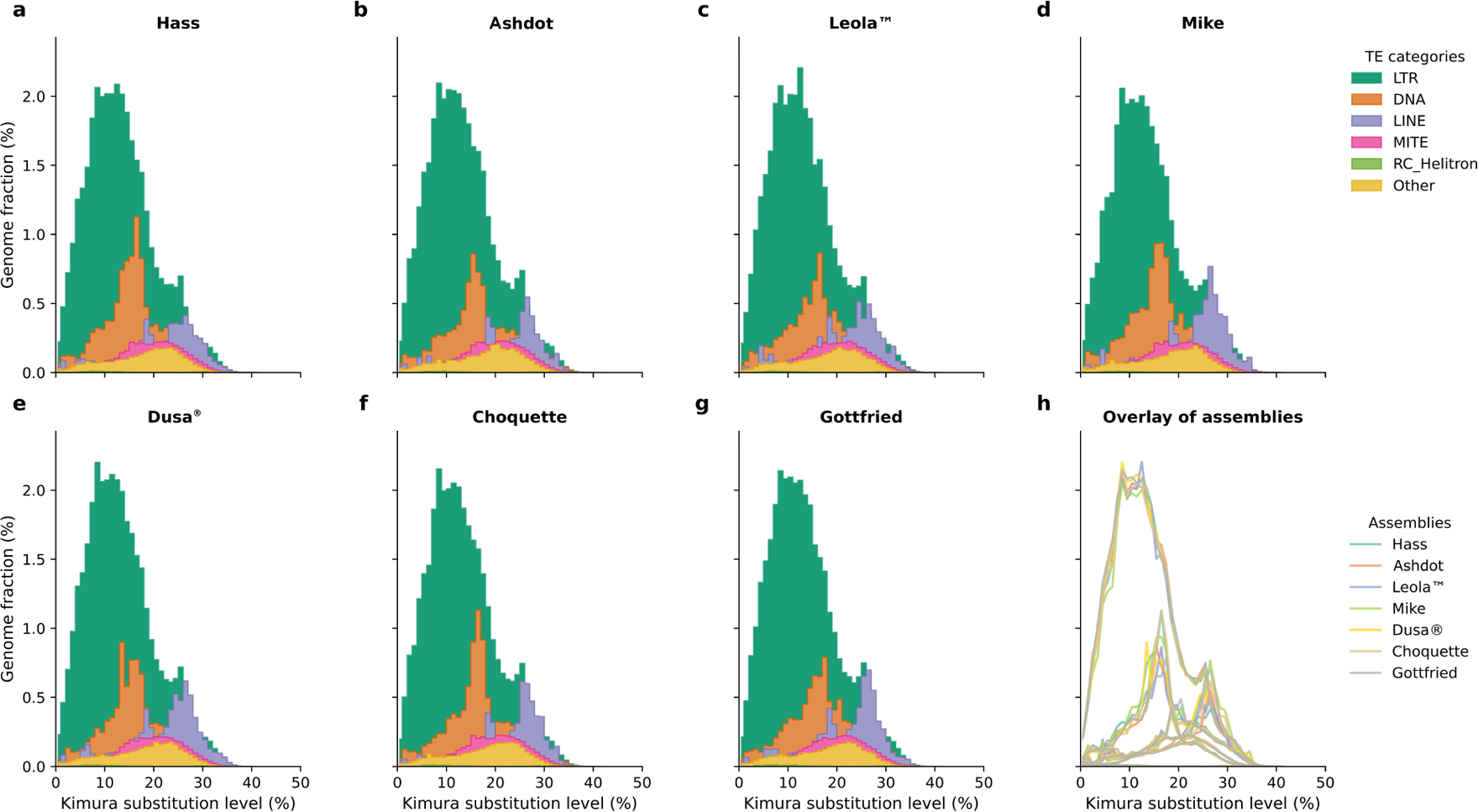
Kimura (percent-substitution) divergence landscapes of broad transposable-element (TE) groups for seven *Persea americana* primary assemblies. **a–g**) Per-assembly TE divergence profiles (x axis = Kimura substitution level, 0–50%; y axis = genome fraction % in each 1% bin) for long terminal repeat (LTR) retrotransposons, DNA transposons, LINEs (long interspersed nuclear elements), MITEs (miniature inverted-repeat transposable elements), RC/Helitron (rolling-circle Helitrons), and Other. **h**) Overlays of the same profiles from all assemblies to facilitate direct comparison. Divergence was calculated from RepeatMasker alignments and binned at 1% resolution; values were normalized to each assembly’s size. The plots highlight a conserved, low-divergence peak (~10% Kimura) dominated by LTRs, with older shoulders indicating ancient TE activity.

### Gene annotation and pangenome content

#### Annotation quality and gene space recovery

Structural annotation of primary assemblies yielded 34,215 (Leola™) to 36,931 (Ashdot) predicted protein-coding genes per accession (Table S4). BUSCO completeness ranged from 97.4% (Hass) to 98.7% (Ashdot). Mean exons per transcript (~5.5) and median gene length (3,953–4,227 bp) were consistent across accessions, and Annotation Edit Distance (AED) distributions indicated well supported models (Fig. S6). Alternate assemblies also captured substantial gene space: BUSCO scores 85.3–97.6% and 31,392 to 34,819 predicted genes. In some cases (e.g. Choquette), the alternate assembly encoded more genes than the primary despite slightly lower BUSCO completeness, illustrating that alternates, though less contiguous, can encode comparable gene sets.

#### Functional annotation

Approximately 86% of gene models received functional descriptions (Table S5). Abundant Pfam domains included those linked to immunity — particularly NLR proteins combining NB-ARC (PF00931; ~12/1k genes) with multiple LRR classes (PF13855, PF00560, PF08263; ~15– 29/1k; Fig. S7, Table S6). These NLR-associated domains ranked alongside protein kinases (PF00069; ~42/1k), pentatricopeptide repeats (PPR; PF01535, PF13041; ~22–25/1k), and F-box proteins (PF00646; ~11–12/1k). Cytochrome P450s (PF00067) and UDP-glucosyltransferases (PF00201) were also prominent.

#### Pangenome partitioning

We partitioned gene space into core, soft-core, shell, and cloud compartments based on orthogroup presence across haplotypes. Because partially phased assemblies capture allelic and structural diversity, each haplotype (primary and alternate) was treated as an independent unit; counts and percentages are reported per haplotype. Across 15 haplotypes (7 accessions x 2 haplotypes plus the WI reference), we identified 38,448 orthogroups: 14,246 core (>95% of haplotypes), 9,005 soft-core (85–94%), 11,393 shell (20–84%), and 3,804 cloud (≤19%; Fig. 2, Fig. S8, Table S7). These corresponded to 17,417–18,155 core genes, 8,107–11,197 soft-core genes, 4,946–6,947 shell genes, and 326–631 cloud genes per assembly.

**Figure 2.**
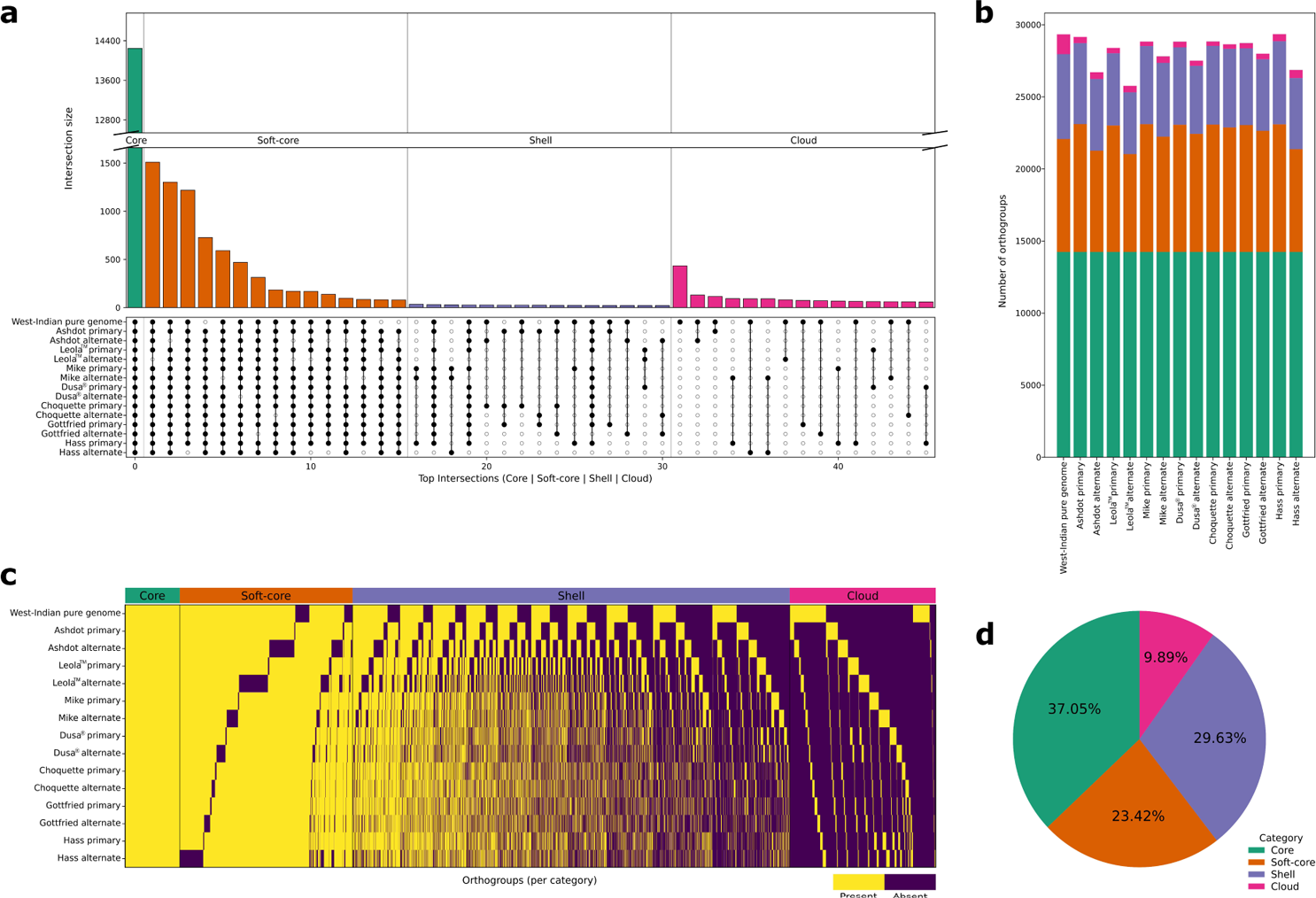
Orthogroup presence–absence variation (PAV) across the avocado pangenome. A binary PAV matrix was derived from orthogroup assignments: a genome is scored present (1) if any of its genes belong to the orthogroup, otherwise absent (0). Colours are consistent across panels by category (Core = green, Soft-core = orange, Shell = purple, Cloud = magenta). Orthogroups were classified by cohort prevalence for n = 15 genomes (West-Indian pure genome; Ashdot, Leola^TM^, Mike, Dusa^®^, Choquette, Gottfried, Hass — each with primary then alternate haplotypes treated separately) using thresholds: Core ≥95% (15/15), Soft-core 85–94% (13–14/15), Shell 20–84% (3–12/15), Cloud <20% (1–2/15). **a)** UpSet-style summary of the most frequent intersection patterns, grouped by category. Bars are coloured by class and the y-axis is split to accommodate the large Core intersection. The dot matrix indicates which genomes contribute to each intersection (row order as above, primary followed by alternate haplotypes). **b)** Per-accession stacked composition of orthogroups present in each genome, coloured by category. **c)** Presence/absence heat map (rows = genomes, columns = orthogroups) sorted by category and prevalence. For legibility, Core and Soft-core are width-compressed; Shell and Cloud are shown at full density. The coloured header strip marks category boundaries. **d)** Pie-chart of the overall pangenome orthogroup category composition.

Relative to the WI reference, 9,107 orthogroups were absent, corresponding to at least 19% novel genes (assuming one gene per orthogroup; Fig. S9), and up to 27.5% novel gene content when accounting for multi-gene orthogroups (13,149 additional genes). Orthogroup accumulation curves indicated an open pangenome (β = 0.109), consistent with continued discovery of novel genes as additional haplotypes are added (Fig. S10). In contrast, sequence-based modelling with Panacus yielded α = 1.138, suggesting that while new orthogroups continue to emerge, broader sequence space is nearing saturation (Fig. S11).

#### Functional enrichment of accessory genome

Gene ontology (GO) enrichment of the accessory genome (shell + cloud) revealed strong over-representation of defence- and stress-related processes. Notably, response to oomycetes, defence response, defence response to Gram-negative bacterium, programmed cell death, and interspecies interactions were enriched (Fig. S12, Table S8). Consistent with immune signalling, calcium-mediated signalling was among the strongest signals, with further enrichment for positive regulation of hydrogen peroxide metabolism and calcineurin–NFAT-related pathways. Specialised metabolism was also prominent, including secondary-metabolite biosynthesis and sesquiterpene pathways, mirrored at the molecular-function level by quercetin glucosyltransferase and sesquiterpene synthase activities. Together with the observed expansions of NLR-associated Pfam domains, these results demonstrate that adaptive immunity and pathogen-response pathways are disproportionately concentrated within the accessory genome.

### Diversity of *NLR* repertoires across accessions

#### NLR identification and classification

*NLR* genes identified with NLRtracker exceeded the 161 *NLRs* previously described in the WI pure accession genome^19^, with 226 (Mike) to 256 (Ashdot) complete *NLR*s (Fig. 3a). Subfamily distributions varied substantially between accessions. Coiled-coil *NLRs* (*CNLs*) dominated; Dusa^®^ encoded the largest set (169) and Hass the smallest (149). Toll/interleukin-1 receptor *NLRs* (*TNLs*) were rare but present in all accessions, with three in Ashdot and Leola™ versus two in all others. Meanwhile, RPW8-like *NLRs* (*RNLs*) were only absent from Choquette.

**Figure 3.**
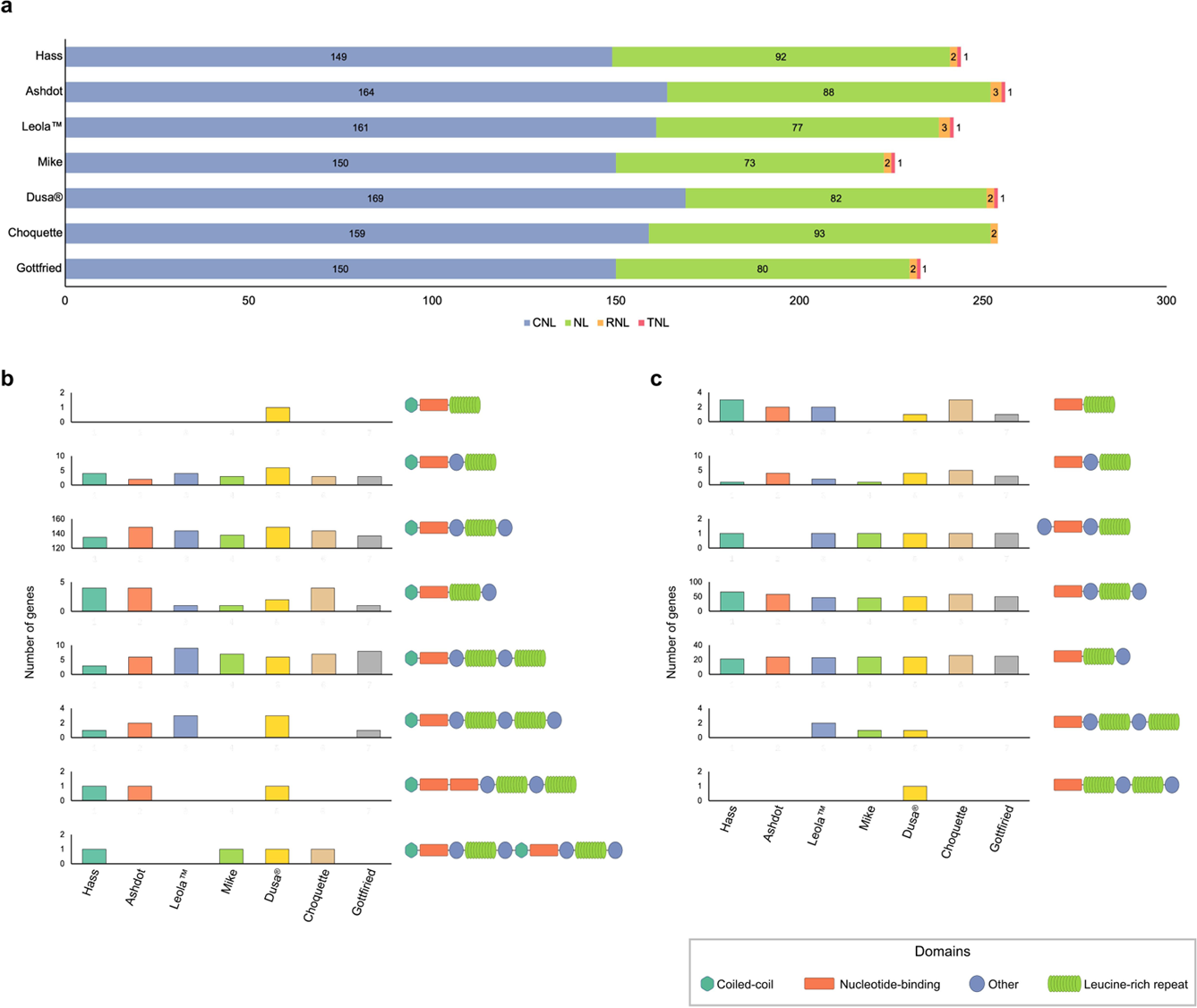
Nucleotide-binding Leucine-rich repeat (*NLR*) gene numbers and domain architectures across seven *Persea americana* accessions. **a)** Total number of genes encoding NLR subfamilies identified in each accession. **b)** Number of CNL-encoding genes partitioned by domain organization. **c)** Number of NLR-type encoding genes carrying alternative domain organizations. Domains were identified with NLRtracker. (C, coiled-coil; N, Nucleotide-binding (NB-ARC); L, Leucine-rich repeat; O, other, integrated or non-canonical domains). Created in BioRender. van den berg, N. (2026) https://BioRender.com/t3s2tsy.

Domain architectures also diverged. Among CNLs, the predominant configuration was C–N– O–L; only a single canonical C–N–L (Dusa^®^) was observed (Fig. 3b). Four accessions (Hass, Mike, Dusa^®^, Choquette) each carried a gene encoding a “C–N–O–L–O–C–N–O–L–O” architecture, suggestive of duplication producing a double NLR within a single protein. All such genes were classified as “cat2” during gene annotation curation, supporting their authenticity. For NLs lacking canonical N-terminal motifs, the most frequent arrangement was N–O–L–O, with Dusa^®^ uniquely carrying N–L–O–L–O (Fig. 3c). Collectively, although copy numbers differ by only a few dozen genes, encoded architectures vary strikingly.

#### Structural variation concentrates at NLR pangenes

Intersecting variants with pangene loci showed markedly elevated SV density within *NLR* windows relative to matched genomic windows (Fig. 4). We observed ~0.35 SV/kb, exceeding the null distribution (Fig. 4a). Single- and multi-nucleotide polymorphisms (SNPs/MNPs) showed a smaller but significant increase of ~15/kb against the null (Fig. 4b). Per-type stratification (Fig. 4c) indicated enrichment driven primarily by inversions and deletions, with insertions also enriched.

**Figure 4.**
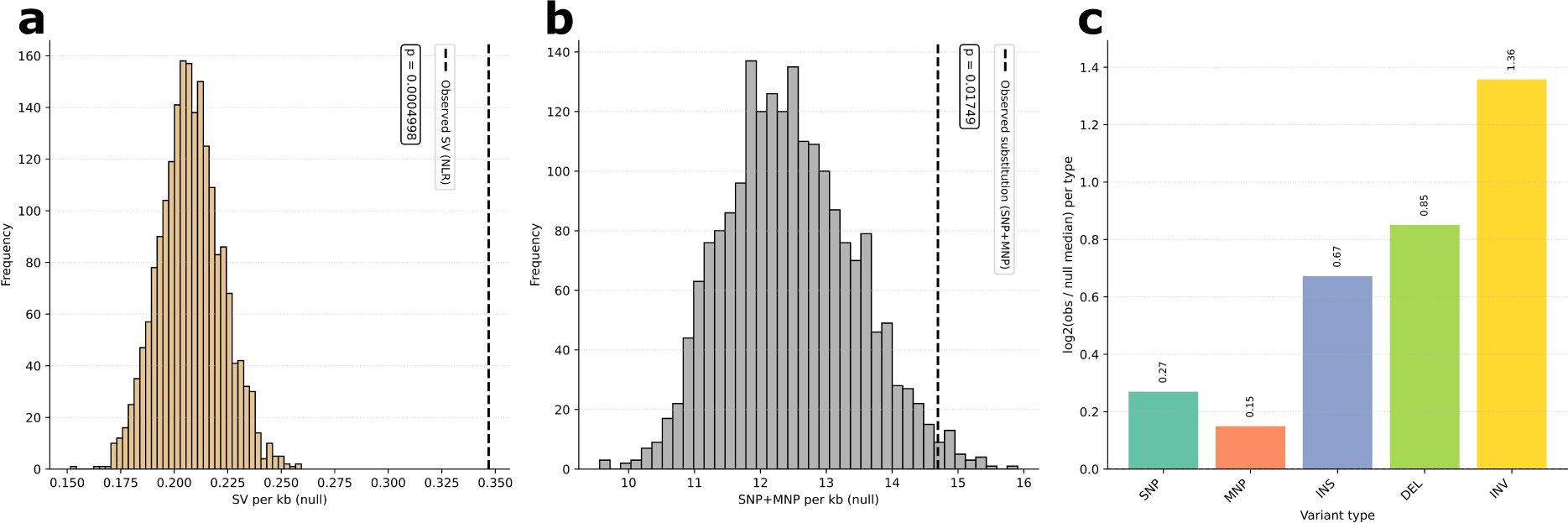
Variation within ± 5 kb of Nucleotide-binding Leucine-rich repeat (*NLR*) loci relative to matched genomic nulls. **a)** The null distribution of structural-variant (SV) density (SVs per kb) in length- and chromosome-matched random windows; the dashed line marks the observed density within ± 5 kb of *NLR* pangenes; *p* from a right-tailed bootstrap (N = 2,000; add-one smoothing). **b)** Null distribution as in panel **a** for substitutions (SNPs + MNPs). Panel **c** displays the log_2_ fold-change of the observed density over the null median for each variant class (log_2_ fold change value above bars); SNP (single nucleotide polymorphism), multi nucleotide polymorphism (MNP), insertion (INS), deletion (DEL), inversion (INV).

Analysis of *NLR* organization within genomic clusters revealed that, on average, 60% of *NLRs* were clustered (Fig. 5a). Gottfried had the fewest clustered *NLRs* (133; 57%), whereas Dusa^®^ had the most (169; 67%). Cluster distribution also varied across chromosomes, with the largest concentrations on chromosome 7; only Ashdot and Dusa^®^ harboured clusters on chromosome 12, and none were detected on chromosome 9 (Fig. 5b). Accessions differed strongly on chromosome 2: Hass carried 32 clustered *NLRs* compared with 16 in Mike. Similarly, Leola™ lacked clusters on chromosome 3, while other accessions averaged ~12 genes.

**Figure 5.**
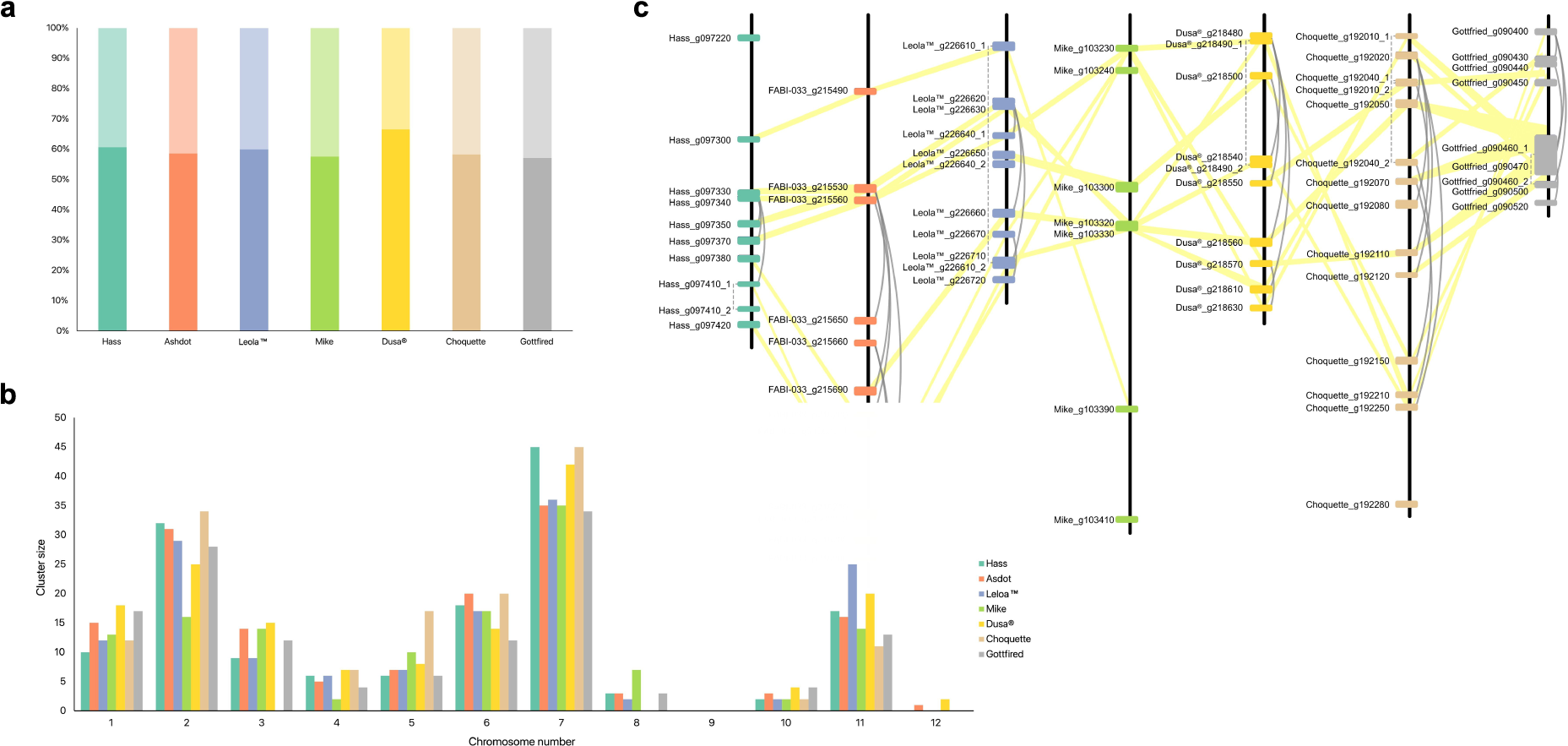
Distribution and organization of clustered Nucleotide-binding Leucine-rich repeat (*NLR*) genes. **a)** Proportion of *NLRs* located within genomic clusters per accession. Dark-coloured segments indicate clustered *NLRs*; transparent segments indicate non-clustered *NLRs*. **b)** Chromosomal distribution of clustered *NLRs*, shown as the total number per chromosome in each accession. **c)** Gene organization of the largest cluster on chromosome 7, compared between all accessions. For each accession, gene identifiers are provided, with grey arcs showing high sequence similarity (>85%) between *NLRs* within a given accession, while yellow connectors highlight *NLRs* with high sequence similarity between different accessions. Grey dash lines are shown for intron regions which span across *NLR* genes. True gene identifiers for different avocado accession are: PeameHass#1 – Hass; PeameRB001#1 – Ashdot; PeameRB002#1 – Leola™; PeameRB003#1 – Mike; PeameRB004#1 – Dusa^®^; PeameRB005#1 – Choquette; PeameRB006#1 – Gottfried.

The largest individual cluster on chromosome 7 comprised 7–14 *NLRs* depending on accession and showed substantial variation in gene order and orientation (Fig. 5c). Comparative sequence analysis indicated multiple, independent duplications within accessions. For example, in Ashdot, two genes (PeameRB003#1_g215650 and PeameRB003#1_g215660) exhibited >98% sequence similarity) to other *NLRs* in the same cluster, but <85% similarity to syntenic *NLRs* in other accessions, suggesting accession-specific duplication. Finally, no significant correlation was detected between *NLR* cluster density and local TE density (r = 0.012, *p* > 0.05), suggesting that recent *NLR* duplications are not primarily driven by TE activity (Fig. S13).

#### Core and accessory NLR repertoires

To test whether *NLRs* skew toward the accessory genome, we quantified *NLR* occupancy across pangenome compartments. On average, 54% of *NLRs* per accession were core (119 in Mike to 140 in Choquette; Fig. 6a), with the remainder consisting ~35% soft-core, ~10% shell, and ~0.4% cloud; Hass and Mike contained no cloud *NLRs*. *NLR* orthogroup accumulation curves indicated an open repertoire (β = 0.147), implying slow but continuing discovery of novel *NLR* orthogroups (Fig. S14).

**Figure 6.**
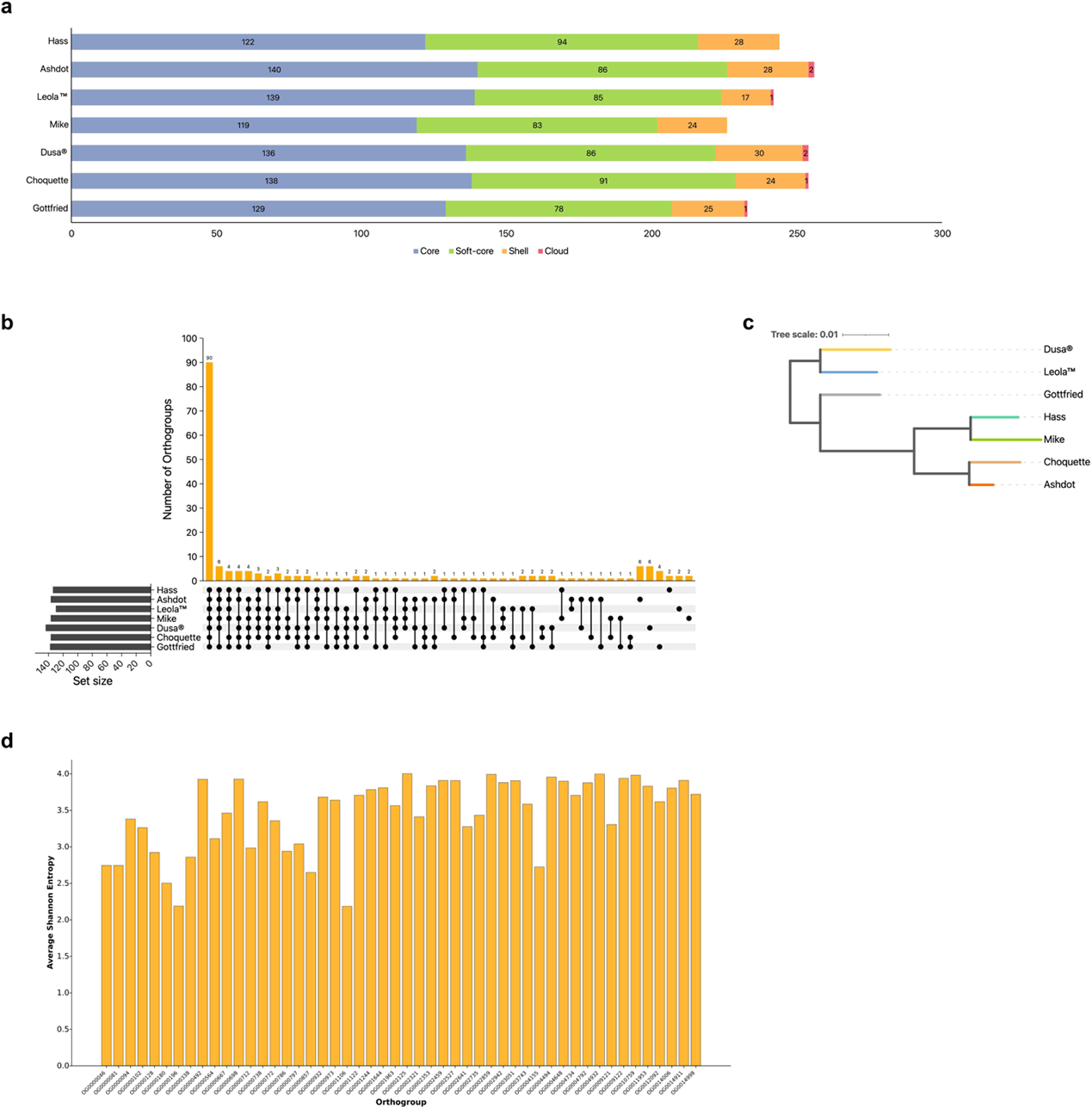
Shared Nucleotide-binding Leucine-rich repeat (*NLR*) genes, functional repertoires, and NLR sequence diversity across avocado accessions. **a)** Number of core, soft-core, shell, and cloud *NLR* genes per accession, classified with GENESPACE. Core genes are present in ≥95% of accessions, soft-core in 85–94%, shell in 20–84%, and cloud in ≤19%. **b)** UpSet plot showing the number of Gene Ontology (GO) terms shared across different combinations of accessions and accession-specific terms. The set size indicates the total number of GO terms identified in each accession. **c)** Multigene species phylogenetic tree inferred with OrthoFinder from the NLR proteome orthogroups. **d)** Average Shannon entropy score per GO term, shown for the top 50 GO terms shared across all accessions. Shannon entropy reflects per-residue amino acid variation within GO term–associated NLRs.

Gene ontology revealed 90 terms shared across all accessions, with the remaining 94 unevenly distributed (Fig. 6b). Each accession thus harbours a partially distinct *NLR* repertoire with limited overlap in accession-specific terms. Phylogenetic analysis of GO term profiles grouped accessions according to genetic relationships: Dusa^®^ and Leola™ clustered together, Hass, Mike, Choquette, and Ashdot formed a second clade, and Gottfried branched separately (Fig. 6c). Average Shannon entropy scores for the top 50 shared GO terms exceeded 2 bits, reflecting substantial amino acid variability (Fig. 6d). Thus, even NLRs that share identical GO annotations across accessions display substantial sequence divergence.

## Discussion

Single reference genomes capture only a portion of a species’ gene space. Crop pangenomes — including tomato^33^, common bean^34^ and apple^9^ — show that reference-only views overlook thousands of genes. By assembling seven *P. americana* accessions spanning all major landraces and hybrid groups and generating near chromosome-scale, partially phased genomes, we reveal an open pangenome: 19–27.5% of genes are absent from the WI reference. Orthogroup accumulation curves indicate continued discovery as additional accessions are added, consistent with other outcrossing species. Avocado’s diversity, shaped by multiple domestication histories^15^, remains unsaturated. Thus, continued germplasm sampling will uncover additional, potentially valuable traits.

Core genes encode essential functions, whereas accessory compartments often harbour adaptive traits^35^. In avocado, shell and cloud genes are strongly enriched for defence- and stress-related responses — programmed cell death, pathogen response, and calcium signalling — with similar patterns reported in crops such as pea^36^ and apple^9^. Thus, a disproportionate share of biotic-stress capacity likely resides in the accessory genome. Underutilised accessions may carry unique resistance, or metabolic traits absent from industry-preferred cultivars and rootstocks, underscoring the importance of conserving broad genetic diversity.

Transposable elements drive genome evolution^12^, yet whether their activity differs among avocado lineages was previously unknown. Across accessions we observed highly conserved repeat content (~59–60%), dominated by LTR retrotransposons, and concordant Kimura divergence profiles. Most LTR activity therefor predates landrace divergence; present-day structural differences largely reflect rearrangements of shared repeats rather than recent bursts, in contrast to maize^37^. This stability provides a robust genomic scaffold while still allowing localized variation.

Linear reference genomes collapse alternative haplotypes, obscuring complex SVs^35^. Our graph-based pangenome embeds SNPs, MNPs, inversions, deletions, and insertions directly in the graph. Meanwhile, orthogroup analyses capture presence/absence variation, revealing diversity often invisible in linear models. Similar graph-based approaches in potato^38^ and wheat^39^ have underlined this strategy for resolving SV-driven trait variation. In avocado, SVs are disproportionately concentrated within *NLR* clusters, highlighting an evolutionary hotspot of immune diversification and paralleling observations in cacao^30^. By representing alternative haplotypes, the graph retains key immune-related SVs rather than collapsing them.

Although overall *NLR* counts are broadly comparable across accessions, structural, organisational, and sequence-level features vary extensively. Domain architectures diverge, including rare canonical or multi-domain configurations that differ between accessions^40^, echoing observations in Arabidopsis^41^ and rice^42^. This indicates that the avocado NLRome is shaped not simply by expansions in gene number but also through the reorganization of protein domain structures and the retention of unusual configurations that may provide functional novelty.

Most *NLRs* occurred in clusters, yet cluster locations, sizes, and arrangements differ; gene order and orientation frequently vary, consistent with independent duplication and rearrangement events^43^. Importantly, these duplication events appear unrelated to TE activity: repeat landscapes are conserved and *NLR* cluster density does not correlate with local TE density, unlike in Arabidopsis^43^. Instead, localized duplication mechanisms seem to have played a greater role in shaping the *NLR* repertoire.

Functional annotation adds resolution: many *NLRs* are shared at the orthogroup level, yet GO terms are unevenly distributed. Even widely shared functions exhibit high Shannon entropy scores, indicating substantial amino acid variation. Reported entropy values exceed those for whole NLR sequences in maize^44^, Arabidopsis and *Brachypodium distachyon*^45^. Thus, proteins with the same annotations are far from uniform across accessions, and diversification within conserved functional classes may be critical for expanding pathogen effector recognition capacity^43^.

Phylogenetic analyses of functional profiles mirrored genetic relationships: related accessions grouped together — consistent with shared domestication and hybridization histories — whereas distant ones followed independent evolutionary trajectories. These patterns emphasize the mosaic nature of the avocado NLRome, where conserved elements coexist with lineage-specific innovations.

Collectively, avocado *NLR* evolution reflects substantial structural heterogeneity with significant functional consequences. Differences in domain organization, cluster dynamics, duplication histories and sequence diversity produce a repertoire richer and more complex than any single genome can capture. Integrating multiple accessions resolves a conserved backbone alongside lineage-specific innovations, providing the most comprehensive view to date.

The avocado pangenome thus reveals a dual architecture: a stable core of genes and conserved repeats, alongside adaptive zones enriched for accessory genes and SVs at *NLR* clusters. This combination provides both resilience and evolutionary flexibility. For breeding, the path forward is to mine accessory diversity and SV variation to improve disease resistance to key pathogens such as *Phytophthora cinnamomi* and *Dematophora necatrix*^46, 47^. Future work should expand sampling to wild relatives, integrate expression and epigenomic data, and experimentally validate candidate loci. Adopting a graph framework ensures that full diversity is represented for association studies and marker design, establishing an integrated avocado pangenome across all major horticultural groups.

## Methods

### Plant material, DNA extraction and sequencing

Six diverse *Persea americana* accessions were selected to capture major genetic backgrounds based on a prior genotyping-by-sequencing (GBS) survey^48^ and in-house genotyping with a 384-marker SNP chip^49^: Ashdot (RB001; West-Indian), Leola™ (RB002; Mexican), Mike (RB003; Guatemalan), Dusa^®^ (RB004; Guatemalan x Mexican), Choquette (RB005; Guatemalan x West-Indian), and Gottfried (RB006; Mexican x West-Indian). To broaden the reference space and benchmark the assembly pipeline, we also incorporated previously generated PacBio HiFi reads from a Hass (Guatemalan x Mexican; Accessions: SRR13510945/6) accession^17^. Young recently expanded leaf flushes were sampled from shaded canopy positions within the germplasm blocks of Westfalia^®^ Fruit Estate, Tzaneen, Limpopo, South Africa.

High-molecular-weight (HMW) DNA was extracted for long-read sequencing using a slightly modified plant HMW DNA extraction protocol^50^. In the modified protocol, (i) DNA was precipitated with chilled isopropanol and incubated overnight at −20°C; (ii) following precipitation, samples were centrifuged at 3,000 x g for 60 minutes at 4°C; (iii) RNase A was added, followed by incubation at 37°C for 15 minutes. The resulting HMW DNA was shipped to Macrogen Inc. (Seoul, South Korea) for SMRTbell library preparation and PacBio Revio sequencing. Libraries were run two samples per flow cell, yielding ~30–40 Gb per sample. Raw HiFi reads were returned as compressed FASTQ files (.fastq.gz) for downstream assembly and analyses.

### Genome assembly

Raw HiFi reads were quality-assessed with FastQC v0.12.1^51^ and reports were aggregated with MultiQC v1.29^52^. Although PacBio HiFi reads are expected to be free of adapter contamination, we verified and removed any residual adapters or artefacts with HiFiAdapterFilt v2.0.0^53^ using the pbadapterfilt.sh wrapper. Read-level summary statistics were generated with fastq-stats (ea-utils^54^), and k-mer spectra were computed with Jellyfish v2.2.10^55^ (k = 20) to produce histograms. The homozygous-coverage peak estimated from these histograms was subsequently used to inform downstream assembly parameters.

Assemblies were generated with Hifiasm v0.25.0^56^ using the computed --hom-cov value and the options -u 1 --telo-m TTTAGGG --telo-s 700 --telo-d 5000 –primary, to recover phased primary and alternate contig sets. The GFA outputs (*.p_ctg.gfa, *.a_ctg.gfa) were converted to FASTA using gfatools gfa2fa v0.5^57^. Resulting primary and alternate FASTA files were carried forward to contaminant removal, repeat annotation, masking and gene prediction.

### Organelle genome assembly and filtering

Organelle genomes (plastid and mitochondrial) were assembled *de novo* from filtered PacBio HiFi reads using OATK v1.0^58^ with the *embryophyta_mito.fam* and *embryophyta_pltd.fam* reference families. Mean genome-wide coverage was estimated by aligning reads to both primary and alternate assemblies with minimap2 v2.29^59^ (map-hifi mode, --secondary=no) followed by depth profiling with samtools v1.22^60^. Organelle-enriched reads were identified using a coverage threshold ≥5x the mean. The resulting sequences were annotated with GeSeq^61^ (Chlorobox) and visualised with OGDRAW^62^.

To remove organelle-associated contigs from the genome assemblies, the mitochondrial and plastid genomes were concatenated into separate reference databases. Nuclear genome contigs were screened against these databases using BLAST*+* v2.16.0^63^ (blastn, E-value [*expected number of chance matches*] cutoff =1e-5, max_target_seqs=1). Contigs with query coverage ≥80%, identity ≥90%, and E-value ≤1e-10 were classified as organelle-associated contigs and removed using seqtk v1.4^64^. Filtering was performed iteratively, first excluding mitochondrial-, then plastid-associated contigs.

### Contaminant identification and filtering

The remaining contigs were taxonomically annotated by protein homology searches against the UniProt Knowledgebase (UniProtKB) SwissProt protein database^65^ (release 2025_01) using DIAMOND v2.1.12^66^ (blastx, ultra-sensitive mode; minimum score 60; ORF ≥30 aa; identity ≥30%; E-value ≤1e-5; top 10 hits per query). The DIAMOND database was built with embedded NCBI taxonomy identifiers^67, 68^. To reduce redundancy, overlapping hits on each contig were collapsed into non-overlapping intervals, retaining only the highest-scoring representative per region. Consensus taxonomy was then assigned per contig by scoring candidate taxonomic identifiers according to unique hit count (2 points), maximum bitscore (1 point), maximum alignment length (0.5 points), and minimum E-value (0.5 points). The taxid with the highest cumulative score was selected as the consensus; in the case of ties, the taxid corresponding to the first best-scoring hit in the filtered BLAST output was chosen.

Contaminant removal was performed with BlobToolKit v4.4.5^69^. For each assembly, BlobToolKit databases were built from organelle-filtered contigs (*blobtools create*), supplemented with coverage profiles from PacBio HiFi read alignments (*minimap2* v2.29; *samtools* v1.22) and taxonomic assignments from DIAMOND consensus hits (*blobtools add*). To ensure correct coverage profiles, BAM files were regenerated after removing organellar contigs. Assemblies were then filtered to retain only contigs assigned to *Viridiplantae* or lacking a taxonomic hit (*blobtools filter* with --query-string “bestsumorder_kingdom-- Inv=Viridiplantae,no-hit”). BlobToolKit outputs included cleaned and contaminant FASTA files, tabular summaries of contig length, GC content, coverage, and phylum-level taxonomy, as well as interactive GC–coverage–taxonomy plots (*blobtools view --plot*) for manual inspection of contamination profiles. The resulting cleaned assembly FASTA files, devoid of organellar and taxonomic contaminants, were used in subsequent analyses.

### Assembly quality assessment and benchmarking

Assembly-level quality was assessed using multiple complementary approaches. Basic assembly statistics, including N50, L50, total assembly length, GC content, and k-mer spectrum profiles, were generated with QUAST v5.3.0^70^. For each accession, raw PacBio HiFi reads were counted with Meryl v1.3^71^ (k=20), and compared against concatenated primary and alternate assemblies in Merqury v1.3^71^ to estimate consensus QV, completeness, and copy-number spectra.

Structural consistency relative to the WI pure reference genome was assessed with minimap2 v2.29 (asm5 preset, –secondary=no). Alignments were converted to sorted BAM files with samtools v1.22, and PAF alignment files were generated with paftools.js v2.29 (sam2paf) for dotplot visualization. Dotplots were inspected with D-GENIES^72^ to evaluate large-scale structural concordance and detect potential misassemblies.

Assembly gene-space completeness was assessed with BUSCO v5.8.3^73^ in genome mode (–m genome) using the embryophyta_odb10 lineage dataset. BUSCO was run with long mode enabled (–long), Augustus species parameter set to *Theobroma cacao* (closest available model), and protein alignment refinement with miniprot. BUSCO full tables were then integrated into BlobToolKit databases for interactive visualization alongside taxonomic and coverage information.

### Annotation of repetitive elements

Repetitive regions in all genome assemblies were annotated using a combination of EDTA v2.2.2^37^, panEDTA, and RepeatMasker v4.1.5^74^. High-confidence coding sequence (CDS) evidence for TE annotation was prepared from the WI pure accession genome (article in preparation) and provided to EDTA to reduce false annotations in gene regions. Repetitive elements were annotated for each assembly with EDTA (--species others --sensitive 1 --anno 1).

A pangenome TE library was then generated by merging individual EDTA annotations with panEDTA, using the high-confidence CDS from the WI pure accession as evidence. The resulting merged library was classified with TEsorter v1.4.7^75^ and subsequently used as a custom repeat database for masking. All assemblies were soft-masked with RepeatMasker, using the curated panEDTA TE library as input.

RepeatMasker outputs were processed in R v4.4.3^76^ with the repeatR package^77^ to quantify and visualize repeat composition. For each assembly we computed (i) the total masked proportion, (ii) TE class composition, and (iii) Kimura substitution profiles by aggregating per-hit divergence (p_sub) into 1% bins from 0–50% and normalizing by assembly size to report genome fraction per bin. For cross-assembly comparisons, subclasses were collapsed into six groups (LTR, DNA, LINE, MITE, RC/Helitron, and Other). For visualization, plots were generated using the ggplot2^78^ and RcolorBrewer^79^ packages.

### RNA-sequencing data acquisition, processing, and normalisation

RNA-seq libraries were retrieved from the NCBI Sequence Read Archive (SRA) database using esearch/efetch (sratools v3.2.1^80^), with the query “Persea americana [Organism] AND illumina [Platform] AND RNA-Seq [Strategy] AND paired [Layout]”. Runs with fewer than 15 million spots were excluded. Sequence data were downloaded with prefetch (sratools) and converted to paired-end FASTQ files using fastq-dump (sra-tools). Adapter trimming and quality filtering (Q ≥30, minimum length ≥50 bp) were performed with fastq-mcf v1.04.807 (ea-utils) using Illumina adapter sequences. Read quality was then assessed with fastq-stats v1.01 (ea-utils), and filtered libraries were retained if they contained ≥15 million reads, mean read length ≥75 bp, mean base quality ≥30, and ≤50% duplication. Runs failing these criteria were discarded. The RNA-seq datasets used are listed in Table S9.

RNA-seq libraries were normalized individually with Trinity v2.15.2^81^ (insilico_read_normalization.pl) using strandedness information curated from metadata and verified with the check_strandedness function in Kallisto v0.44.0^82^. Libraries were normalized to a maximum coverage of 100x to (i) prevent any single library from dominating downstream analyses, (ii) balance contributions across samples, and (iii) reduce overall computational load. Libraries were processed separately to preserve sample integrity and maintain a clean output structure for subsequent alignment.

### *S*tructural annotation

RNA-seq alignments were performed using STAR v2.7.11b^83^ with a two-pass approach on individually normalized libraries. STAR genome indices were built for both primary and alternate assemblies, and resulting alignments were sorted by coordinate with non-canonical splice junctions removed. For gene prediction, we applied two complimentary approaches: BRAKER3 v3.0.8^84^, which integrated RNA-seq alignments and protein homology from OrthoDB v12 using the *Viridiplantae* partition^85^, and Helixer v0.3.5^86^ which performed *ab initio* prediction using the land_plant model. Annotation quality was assessed with AED scores generated with InGenAnnot v0.0.15^87^. AED values were calculated relative to RNA-seq transcript assemblies (StringTie2 v2.2.1^88^) and protein alignments against the proteomes of closely related species and the UniProtKB SwissProt reviewed protein database (miniprot v0.17^89^; Table S10), producing both transcript- and protein-based AED metrics.

Because Helixer predictions do not account for TE, Helixer gene models were filtered with an in-house Python script (Filter_repeat_region_genes.py, available: https://github.com/RobBacker/Genome_assembly_and_annotation_tools). The script calculates CDS-level overlap with repeat regions (RepeatMasker annotation file), merges CDS and repeat intervals to avoid double-counting, and removes models with >25% CDS overlap with repeats unless rescued by transcript-based AED (aed_ev_tr < 0.75). BRAKER3 and Helixer annotations were subsequently merged into a unified gene set with AGAT v1.4.2^90^ (agat_sp_complement_annotations.pl), setting the BRAKER3 annotation as the reference. RNA-seq-based transcript evidence was then clustered, ranked, and UTRs were added using InGenAnnot (clusterize, isoform_ranking, and utr_refine), guided by strandedness-aware BAM inputs.

### Gene model curation

Gene model quality was evaluated with PSAURON v1.0.6^91^ using default settings. To guide automated curation, an in-house Python script (aed_psauron_curation.py) combined AED scores with support inferred from PSAURON. Each transcript was assigned to one of six categories (cat1–cat6) ranging from high confidence (cat1: strong transcript and protein support) to *ab initio* only (cat6: no external support). Categories cat1–cat4 were automatically retained, cat5–cat6 were retained only if PSAURON support was detected. This produced an annotation file with per-transcript support categories and PSAURON tags. Diagnostic plots of transcript vs. protein AED distributions were generated to visualize category assignments (curated_aed_psauron_scatter.py).

Curated annotations were then filtered with AGAT utilities to remove low-confidence or pseudogenic models, and summary statistics were generated. For downstream benchmarking, the longest isoform per gene was extracted (agat_sp_keep_longest_isoform.pl) and assessed with BUSCO.

Annotation of TE-related genes was performed using DeTEnGA^92^, which integrates sequence similarity searches against curated TE protein families. For each assembly (primary and alternate), DeTEnGA was run via the GAQET2 wrapper with the plant-specific REXdb database (“rexdb-plant”), using default parameters. The curated GFF3 annotation file served as input, alongside the repeat-masked genome assembly. DeTEnGA produced TE annotation summaries for assembly, which were subsequently used to tag candidate TE-related models using a custom Python script (tag_detenga.py).

### Functional annotation

Functional annotation of predicted proteins was performed using EnTAP v2.3.0^93^. Longest isoform CDS from both primary and alternate assemblies were used as input. EnTAP was run with a *Viridiplantae* taxonomic scope, designating bacterial, fungal, insect, and viral sequences as contaminants. Reference databases included RefSeq Plant and UniProt SwissProt, while domain-based annotation employed InterProScan v5.74-105.0^94^ (databases: Pfam, Panther, SMART, Superfamily, Gene3D, PrositeProfiles, and PRINTS). EnTAP was executed with ontology assignment enabled (GO and KEGG). Functional annotations were parsed and integrated with structural annotation features using AGAT (agat_sp_manage_functional_annotations.pl). Pfam domain summaries were generated from the EnTAP tables using a custom script (entap_pfam_qc.py), which collapses isoforms, counts unique gene–Pfam pairs, and normalizes counts to per-1,000 genes.

### Pangenome graph assembly and assessment

Assemblies and annotations were reformatted to the PanSN-spec naming scheme, with headers encoded as *sample#haplotype#contigID* (haplotypes: 1 = primary, 2 = alternate, 0 = reference). Gene models were updated accordingly, and multi-isoform loci were collapsed to the longest isoform using AGAT.

A whole-genome alignment–based pangenome graph was then constructed with Minigraph-Cactus v8.1.0b1^95^, using the WI pure accession as the reference backbone and incorporating both haplotypes of each assembly in order of k-mer completeness. Runs were performed with the --lastTrain option and produced multiple formats (--odgi, --gfa full clip, --gbz, --vcf, --giraffe, --viz, --vcfwave).

Graph growth statistics were calculated with Panacus v0.3.3^96^ from GFA walk records at the bp level, using quorum thresholds (-q 1,0.5,0.1) and coverage levels (-l 1,3,6,7), both with and without sample merging (-S). Curves were visualized with panacus-visualize.

Orthology and synteny were resolved with GENESPACE v1.3.1^97^, using BED files derived from GFF3 annotations and peptide FASTAs (longest isoform per gene). Primary and alternate haplotypes were treated as independent genomes to maximise discovery of allelic variation across accessions, with the WI reference processed in parallel. Orthogroup presence/absence was analysed with a custom workflow (pangenome_pav_support.py), built around OrthoFinder v2.5.5^98^ outputs from GENESPACE. This produced: (i) a binary presence/absence variation (PAV) matrix, (ii) an orthogroup classification table, and (iii) per-assembly gene-level support matrices from curated GFF3 annotations. Category cutoffs were set to match whole-haplotype counts for our dataset (n = 15): Core ≥95% (15/15), Soft-core 85–94% (13–14/15), Shell 20– 84% (3–12/15), Cloud <20% (1–2/15). Cloud orthogroups were further divided into “Cloud-supported” and “Cloud-unsupported” based on AED curation tags. Plots were generated from the binary orthogroup PAV matrix with a custom Python script (upset_orthogroups.py), which: (i) selects the top 15 intersections from each non-core category for the UpSet-style panel (split y-axis), (ii) draws per-accession stacked compositions, and (iii) renders a presence/absence heat map sorted by category and prevalence.

Pangenome accessory genes (Cloud and Shell) were tested for GO enrichment with topGO v2.58.0^99^ (classic Fisher’s exact test; BP, MF, CC ontologies), restricting terms to the *Viridiplantae* lineage using the GO taxon constraints ontology (go_taxon_constraints.owl^100^). Statistical significance was adjusted using the Benjamini–Hochberg false discovery rate (FDR) procedure. Results were simplified with clusterProfiler v4.14.6^101^ (semantic similarity, cutoff = 0.7) and visualized as dotplots with enrichplot.

Pan- and core-genome accumulation curves were derived from the binary orthogroup PAV matrix using 1,000 random genome order permutations. At each increment, mean, median, and 95% confidence intervals of pan- and core-genome sizes were computed. The pangenome was modeled by Heaps’ law (Pan(n) = S·n^β) using log–log regression, and the core-genome with a Tettelin-style exponential decay model (Core(n) = Ω + k·e^−n/τ). Model parameters and statistics are reported in Table S11. Curves, model fits, and inset log–log regressions were visualized in R v4.4 with ggplot2.

### Pangenome graph structural variant analysis

From each assembly, the longest gene model per locus was mapped to the WI reference in the Minigraph-Cactus graph using halLiftover v2.2^102^. Annotations were standardised with awk and converted to BED with bedtools v2.31.1^103^, after which each mapped gene was reduced to a single span. Pangene loci were defined as regions were these spans overlapped on the same strand across assemblies; with locus coordinates taken as the union of all member spans. From this, we generated a binary PAV matrix and a pangene–gene ID map.

Graph variants were obtained from the Minigraph-Cactus WAVE VCF and processed with bcftools v1.22^60^ to extract SVs (bcftools view -i ‘INFO/INV=1 || INFO/LEN<=−50 || INFO/LEN>=50’) and substitutions (SNPs and MNPs; bcftools view -i ‘INFO/TYPE=“snp” || INFO/TYPE=“mnp”’). Records were exported to BED8 (one line per alternate allele) with TYPE and LEN fields. Pangene loci were then intersected with these variant BEDs using bedtools to obtain per-locus overlap tables and type-stratified summaries for enrichment analyses and figures. This workflow is generalised in the bash script, extract_pan_gene_loci.sh. Variation in and around *NLRs* was assessed by comparing observed variant density within ±5 kb windows around *NLR*-associated pangene loci to a length- and chromosome-matched bootstrap null. For each *NLR* window, a random genomic window of identical length was sampled, with any overlap with the observed *NLR* set excluded. The bootstrap null was generated from 2,000 replicates and, per replicate, overlaps with SVs, SNPs and MNPs were counted with bedtools intersect, reporting densities as variants per kb and stratifying by type. Right-tail significance used add-one smoothing: p = (1 + #(null ≥ obs)) / (N + 1). SV and SNP+MNP bootstraps shared random seeds for directly comparable nulls. Analyses and plotting were performed with a custom Python script, sv_enrichment.py.

### NLR data analysis

NLR-encoding genes were identified from predicted protein sequences using NLRtracker^40^. Genes were classified as complete *NLRs* if they contained at least a central Nucleotide-binding (NB-ARC) domain and a C-terminal Leucine-rich repeat (LRR). N-terminal domains were further categorized as coiled-coil (CC), Toll/interleukin-1 receptor (TIR), or other (O) non-canonical integrated domains (classified as “Other” by NLRtracker). Domain composition schematics were generated with BioRender, and summary statistics were visualized in Microsoft Excel.To assess genomic clustering, GFF3 annotations were used to map *NLR* positions. A cluster was defined as ≥2 *NLR* genes located within 250 kb of one another with <5 non-*NLR* genes between neighbouring *NLRs*. Chromosomal assignment of clusters was determined using the contig–chromosome alignments from whole-genome scaffolding.

Detailed analysis of chromosome 7 clusters was performed with BLAST+ v2.13.0. Genes were considered sequence-similar if they shared >85% nucleotide identity. Local duplication patterns and gene orientations were visualised using custom Python scripts (chromosome_viz.py). To investigate relationships between *NLRs* and TEs, we developed a custom Python script (gff3_density_analyzer.py). The script parsed GFF3 annotation files (for main gene annotations, *NLRs*, and TEs) and determined chromosome boundaries from annotated genes. *NLR* and TE densities were then calculated in non-overlapping sliding 50 kb windows, with values smoothed using a Gaussian filter. A scatter plot of paired *NLR* and TE densities across windows were generated, and the Pearson correlation coefficient was computed to quantify the association between *NLR* and TE distributions. *NLRs* were assigned to core, soft-core, shell, or cloud compartments using orthogroup classifications from GENESPACE v0.9.0 in combination with functional annotation derived from GO analysis. GO terms were also used to construct an UpSet plot, summarizing shared and accession-specific functional annotations. Classification thresholds followed the same definitions applied to the whole-genome pangenome (core ≥95% of haplotypes; soft-core 85–94%; shell 20–84%; cloud ≤19%).

Orthogroups of NLRs from all accessions were further analysed with OrthoFinder v2.5.5 to infer a species-level phylogeny. The analysis was automated with a custom Python pipeline (orthofinder_phylogeny.py), which validated input FASTA files, executed OrthoFinder with default parameters, and produced species trees. Where applicable, IQ-tree was used for tree inference. The resulting tree was visualized using the iTOL webtool.

To assess amino acid diversity within functional groups, protein sequences of NLRs annotated with the same GO term were aligned using MUSCLE v5.3. Shannon entropy was calculated for each alignment using ShannonEnt (available at: https://github.com/wldolan/shannon-entropy), and average entropy scores per GO term were computed. These values were used for graph construction using a custom Python script (orthogroup_pipeline.sh), and to compare sequence variability across shared functional categories.

## Data Availability

Raw long-read sequencing data have been deposited in the **Sequence Read Archive (SRA)** under BioProject PRJNA1331494 (sample accessions: **SRR35509374 – SRR35509379**). Nuclear genome assemblies are available in **GenBank** under accessions **JBRKCI000000000**, **JBRKCJ000000000**, **JBRKCK000000000**, **JBRKCL000000000**, **JBRKCM000000000**, **JBRKCN000000000**, **JBRKCO000000000**, **JBRKCP000000000**, **JBRKCR000000000**, **JBRKCS000000000**, **JBRKCT000000000**, **JBRKCU000000000.**

## Code Availability

Custom Python scripts referenced throughout this article are available on GitHub at https://github.com/RobBacker/Genome_assembly_and_annotation_tools.

## Acknowledgements

We thank the Avocado Genome Consortium for providing pre-publication access to genome resources. We gratefully acknowledge Westfalia^®^ Fruit Estate for providing access to their germplasm and authorizing its research use, and ZZ2 for authorizing the research use of Ashdot material. Institutional support was provided by the University of Pretoria and the Forestry and Agricultural Biotechnology Institute (FABI). This research was funded by the Hans Merensky Legacy Foundation.

## Author Contributions

NB, AC, AV, AB and RB conceived the study. AV optimized DNA extraction and performed all wet-lab experiments. RB generated genome assemblies, developed analysis scripts and conducted primary analyses; AC performed fine-grained NLR analyses. AB provided initial conceptual guidance and mentoring. NB provided resources, supervision, project administration and funding acquisition. RB and AC wrote the original draft. All authors (RB, AC, AV, AB, NB) reviewed and edited the manuscript.

## Supplementary Information

Supplementary materials can be found in Supplementary_Tables.xlsx and Supplementary_Figures.docx.

