## Supplementary_Figures for "The avocado pangenome reveals dynamic clustering and lineage-specific diversity of *NLR* genes"

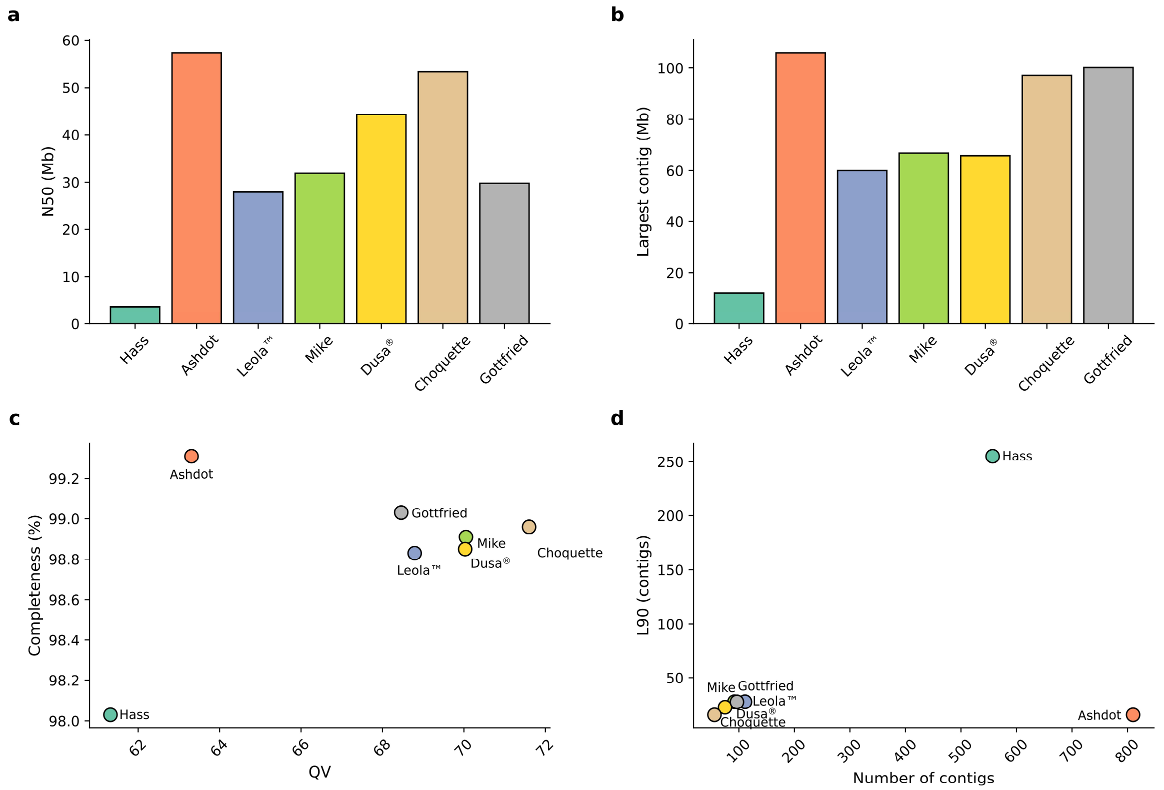


**Supplementary Figure 1. Assembly contiguity and quality across seven *Persea americana* accessions.** **a**) Contig N50 (Mb). **b**) Length of the largest contig (Mb). **c**) Merqury completeness (%) plotted against QV. **d**) Number of contigs versus L90 (number of contigs needed to cover 90% of the assembly length). Bars and points are coloured consistently by accession (Hass, Ashdot, Leola$^{TM}$, Mike, Dusa®, Choquette, Gottfried). Mb, megabases; QV, Merqury k-mer-based consensus quality value.

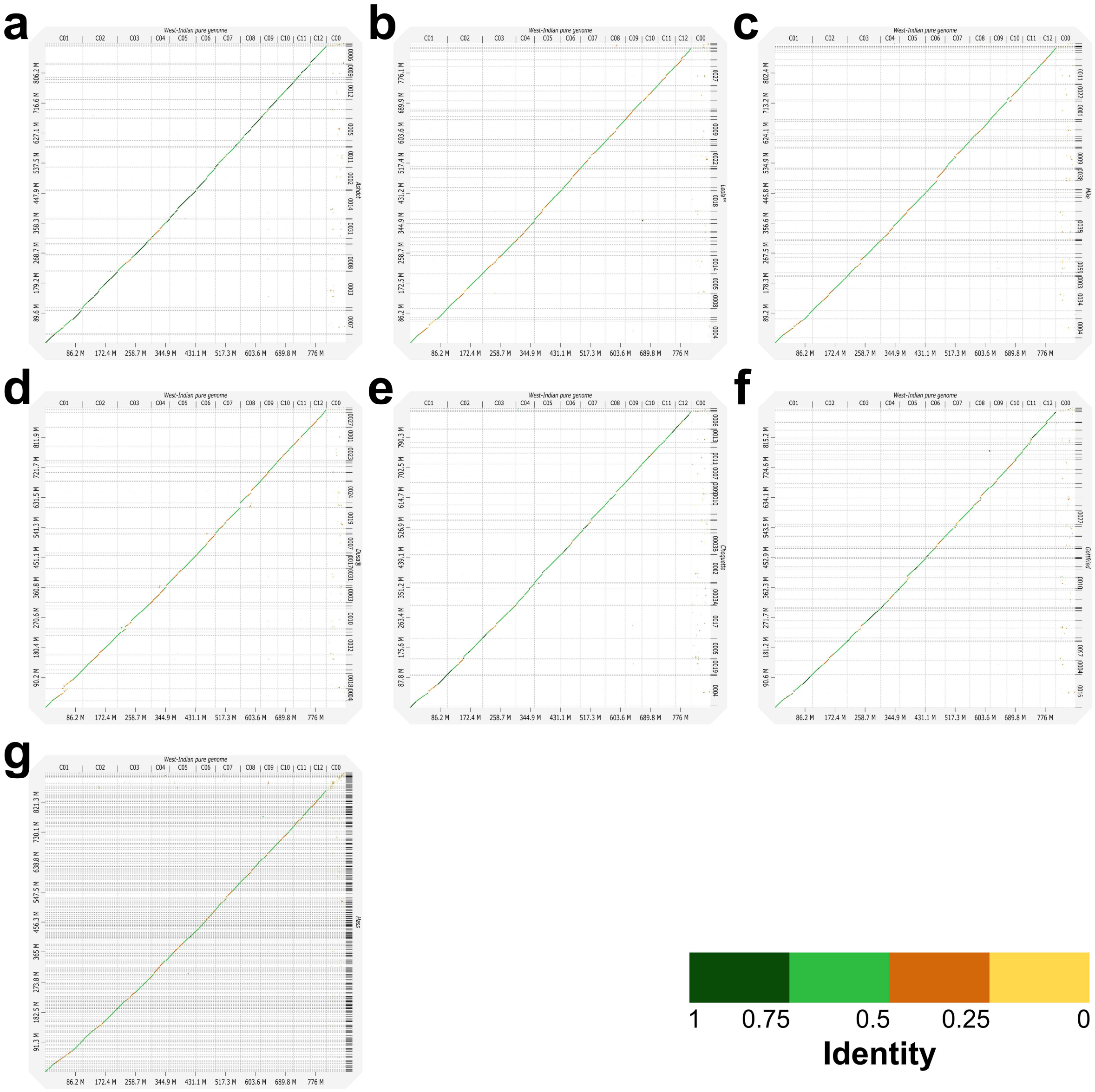


**Supplementary Figure 2. Whole-genome dot plots of seven *Persea americana* assemblies against the West-Indian pure reference.** Pairwise alignments (minimap2 v2.29[1], asm5; unique hits) are shown as points coloured by sequence identity (green = higher, yellow = lower; scale at bottom). The x-axis shows reference chromosomes C01–C12 (C00 = unplaced); the y-axis lists the primary scaffolds of each accession. **a**) Ashdot, **b**) Leola™, **c**) Mike, **d**) Dusa®, **e**) Choquette, **f**) Gottfried, **g**) Hass. A near-continuous 1:1 diagonal indicates chromosome-scale collinearity, whereas small off-diagonal signals denote local rearrangements, repeat-associated matches, or assembly breaks. Plots were generated from PAF alignments with D-GENIES[2].

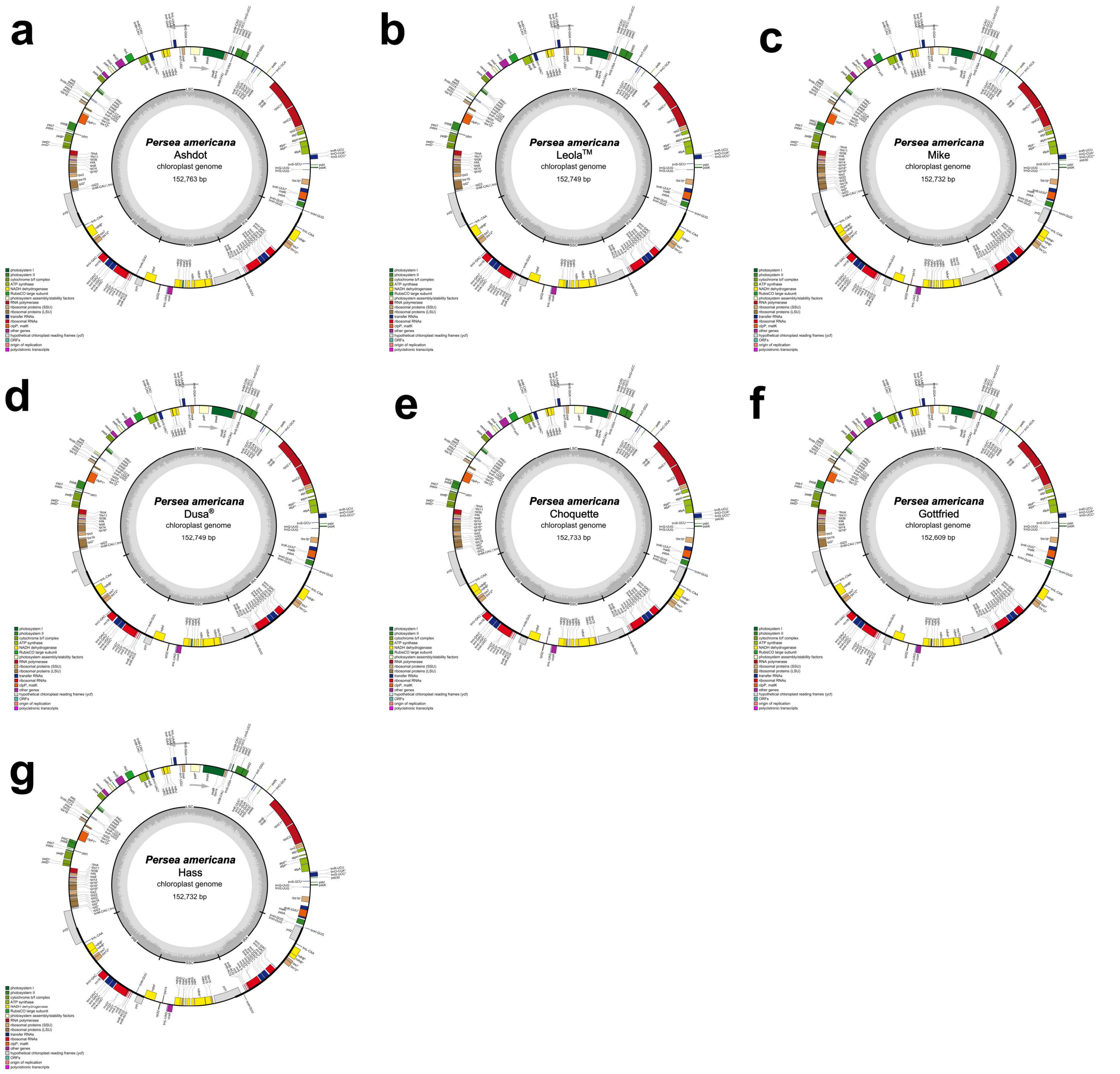


**Supplementary Figure 3. Chloroplast genomes of seven *Persea americana* accessions.** Circular maps of the plastomes for **a**) Ashdot (152,763 bp), **b**) Leola™ (152,749 bp), **c**) Mike (152,732 bp), **d**) Dusa® (152,749 bp), **e**) Choquette (152,733 bp), **f**) Gottfried (152,609 bp), and **g**) Hass (152,732 bp). Genes are coloured by functional class (legend inset). Boundaries of the two inverted repeats ($IR_A$, $IR_B$), the large single-copy (LSC), and the small single-copy (SSC) regions are indicated; genes drawn on the inside/outside of each ring are transcribed clockwise/counter-clockwise, respectively. The inner grey plot shows GC content. Annotations were generated with GeSeq[3] and rendered with OGDRAW[4].

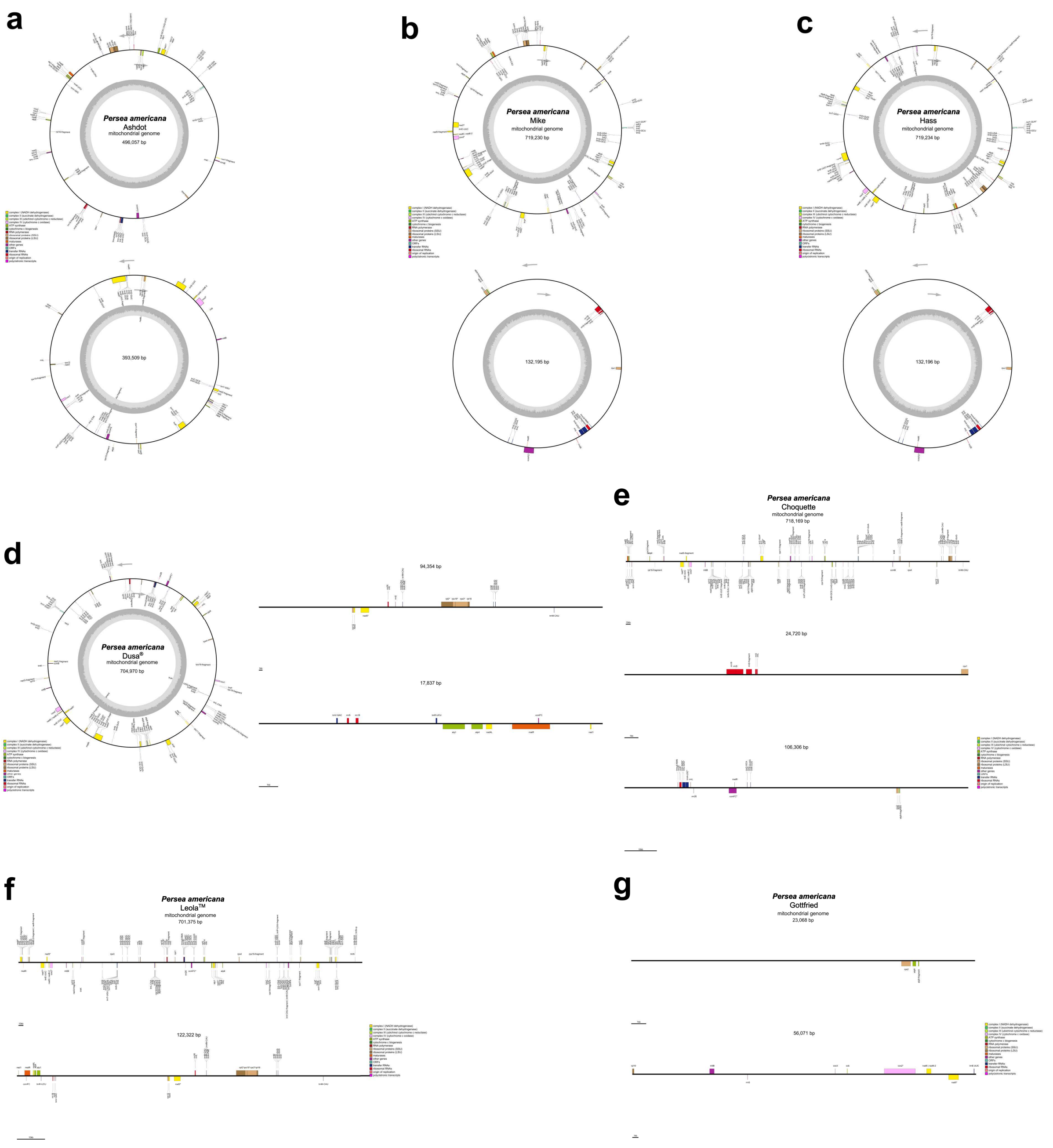


**Supplementary Figure 4. Mitochondrial genomes of seven *Persea americana* accessions.** Maps of assembled mitochondrial contigs with genes coloured by functional class (as in Supplementary Fig. S2) and inner grey rings showing GC content. **a**) Ashdot, two circular contigs (496,057 bp and 393,509 bp); **b**) Mike, two circular contigs (719,230 bp and 132,175 bp); **c**) Hass, two circular contigs (719,234 bp and 132,196 bp); **d**) Dusa®, one circular contig (704,970 bp) plus two linear contigs (94,354 bp and 17,837 bp); **e**) Choquette, three linear contigs (718,169 bp, 106,306 bp, 24,720 bp); **f**) Leola™, two linear contigs (701,375 bp and 122,322 bp); **g**) Gottfried, incomplete assembly represented by two short linear contigs (56,071 bp and 23,068 bp). Annotations were generated with

GeSeq[3] and rendered with OGDRAW[4]. (Note that “circular”/“linear” reflects the assembly representation of contigs rather than definitive *in vivo* topology.)

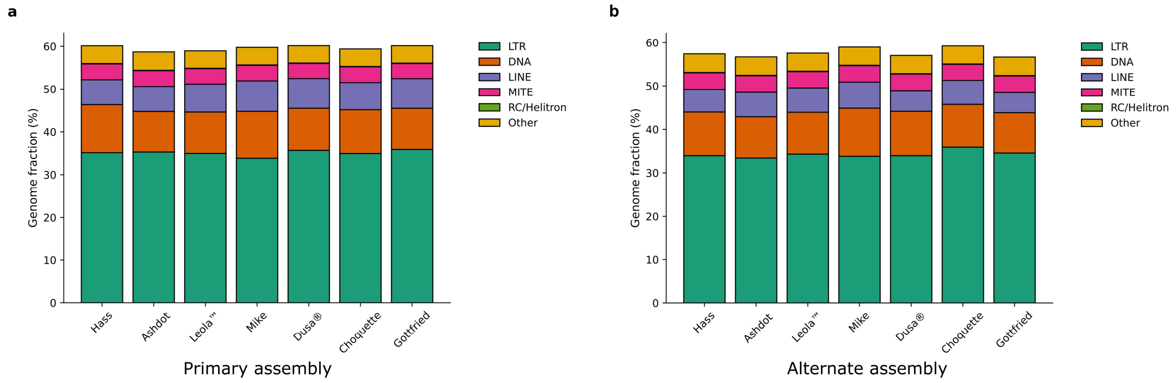


**Supplementary Figure 5. Genome-wide transposable-element composition across seven *Persea americana* primary and alternate assemblies.** Stacked barplot showing the fraction of each genome (%) occupied by broad TE categories (LTR = long terminal repeat retrotransposons; DNA = DNA transposons; LINE = long interspersed nuclear elements; MITE = miniature inverted-repeat transposable elements; RC/Helitron = rolling-circle Helitrons; Other = low-complexity, simple repeats and unclassified). Values are based on repeat annotations obtained with EDTA/panEDTA v2.2.2[5] and RepeatMasker v4.1.5[6] using the curated pan-TE library (see Methods and Table S3 for full breakdown by subclass). Colours correspond to TE categories as plotted. The stacked bars show that repeats comprise ~59–60% of each primary assembly (**a**), and ~56-59% of alternate assemblies (**b**), with LTR elements the dominant class and an appreciable fraction remaining unclassified (LTR/unknown).

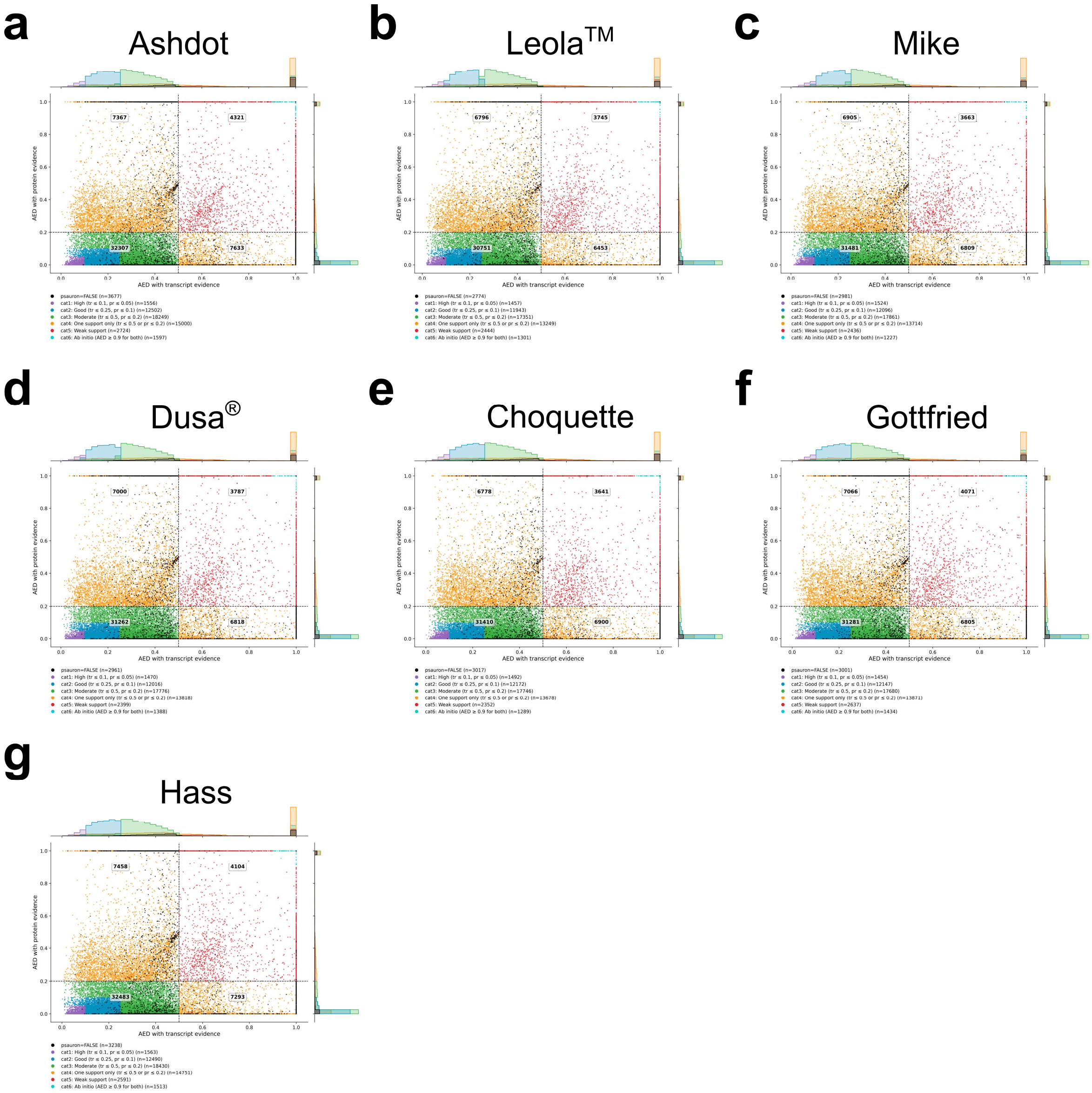


**Supplementary Figure 6. Annotation Edit Distance (AED) summaries for the primary assemblies of seven *Persea americana* accessions.** **a**) Ashdot, **b**) Leola™, **c**) Mike, **d**) Dusa®, **e**) Choquette, **f**) Gottfried, **g**) Hass. Each panel plots RNA-seq–based AED on the x-axis and protein-homology AED on the y-axis (0 = perfect agreement, 1 = no supporting evidence). Points represent gene models categorised by AED scores: cat1 high confidence (transcript AED ≤ 0.10 and protein AED ≤ 0.05; purple), cat2 good (≤ 0.25 and ≤ 0.10; blue), cat3 moderate (≤ 0.50 and ≤ 0.20; green), cat4 one evidence type only (transcript ≤ 0.50 **or** protein ≤ 0.20; orange), cat5 weak (above those bounds but below ab-initio; red), and cat6 ab-initio only (both ≥ 0.90; turquoise). Models classified as ‘FALSE’ by PSAURON v1.0.6[7], are coloured in black. Dashed lines at transcript AED = 0.50 and protein AED = 0.20 mark the “moderate” thresholds; numbers printed in each quadrant report model counts. Marginal histograms summarize AED distributions along each axis.

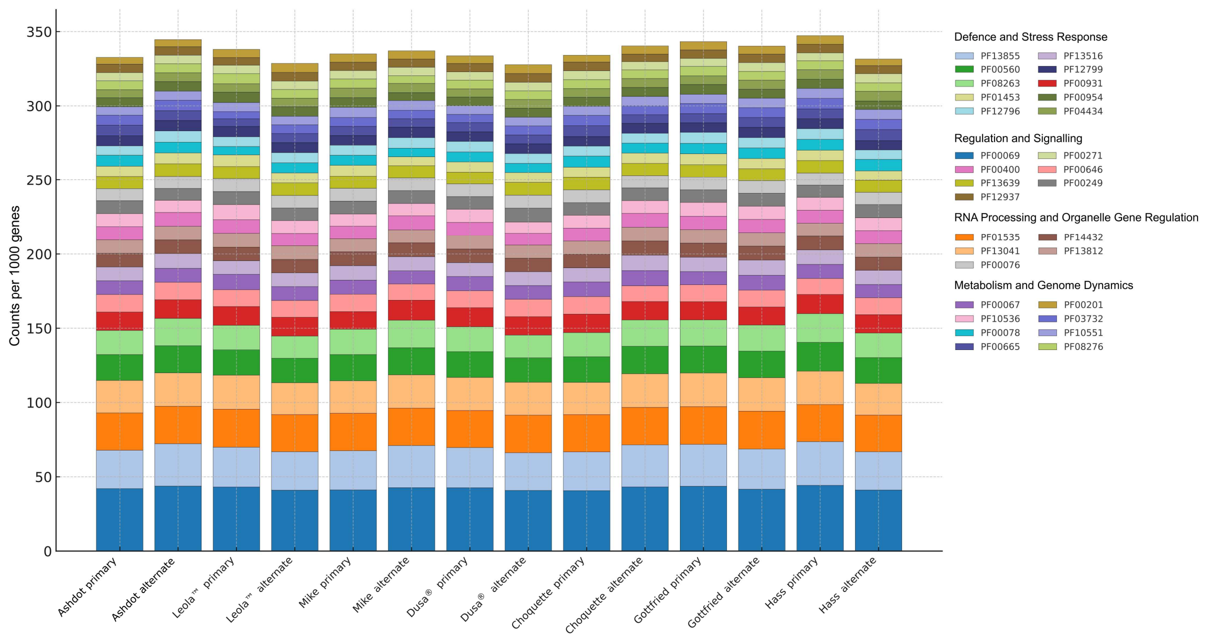


**Supplementary Figure 7. Most abundant Pfam protein domains across the seven *Persea americana* assemblies.** Stacked bars show domain counts per 1 000 genes (isoform-collapsed; each gene–Pfam pair counted once) for the top, recurrent domains, with both primary and alternate haplotypes shown for each accession. Colours correspond to individual Pfam accessions and are grouped in the legend into four functional classes: Defence & Stress Response (PF13855 leucine rich repeat (LRR), PF00560 LRR, PF08263 leucine rich repeat N-terminal, PF00931 NB-ARC, PF13516 LRR, PF12799 LRR, PF01453 D-mannose-binding lectin, PF00954 S-locus glycoprotein, PF12796 Ankyrin repeats, PF04434 SWIM zinc finger), Regulation & Signalling (PF00069 protein kinase, PF00400 WD/G-beta, PF13639 RING finger, PF12937 F-box-like, PF00646 F-box, PF00249 Myb-like DNA-binding, PF00271 helicase C-terminal), RNA Processing & Organelle Gene Regulation (PF01535 pentatricopeptide repeat (PPR), PF13041 PPR family, PF13812 PPR domain, PF14432 DYW deaminase, PF00076 RNA recognition motif), and Metabolism & Genome Dynamics (PF00067 cytochrome P450, PF10536 plant mobile domain, PF00078 reverse transcriptase, PF03732 retrotransposon gag, PF00665 integrase core, PF10551 MULE transposase, PF00201 UDP-glucuronosyl/UDP-glucosyl transferase, PF08276 PAN-like).

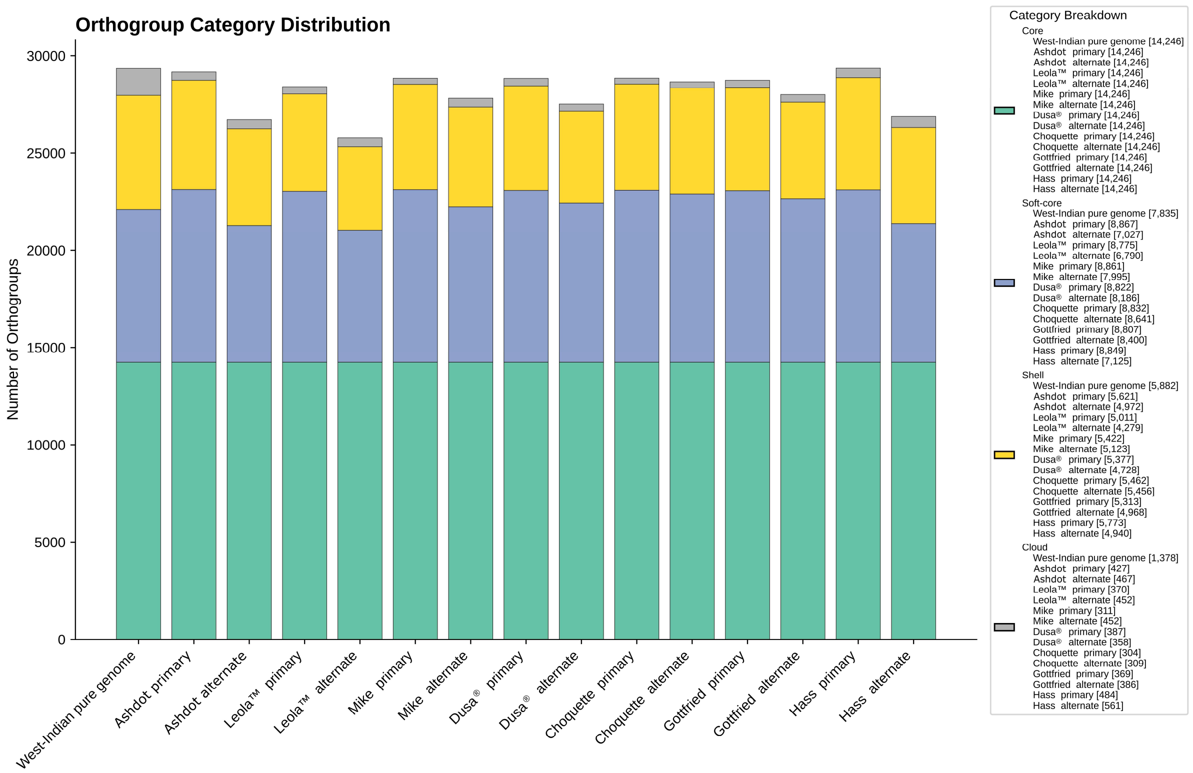


**Supplementary Figure 8. Orthogroup category distribution across *Persea americana* assemblies.** Stacked bars show, for each assembly (left–right: West-Indian pure accession genome; Ashdot, Leola$^{TM}$, Mike, Dusa®, Choquette, Gottfried, and Hass — primary and alternate where available), the number of orthogroups present in each assembly are binned by pangenome class. Classes were defined by across-haplotype frequency: Core (> 95% of haplotypes; turquoise), Soft-core (85 – 94%; blue), Shell (20 – 84%; yellow), and Cloud (≤ 19%; grey). Orthogroup counts per assembly are displayed in the legend on the right. Orthogroups were inferred with OrthoFinder v2.5.5[8] (run within GENESPACE v1.3.1[9]) using longest-isoform peptides, and presence/absence classification was generated with the pangenome workflow (pangenome_pav_support.py). The y-axis reports the number of orthogroups per category for each assembly.

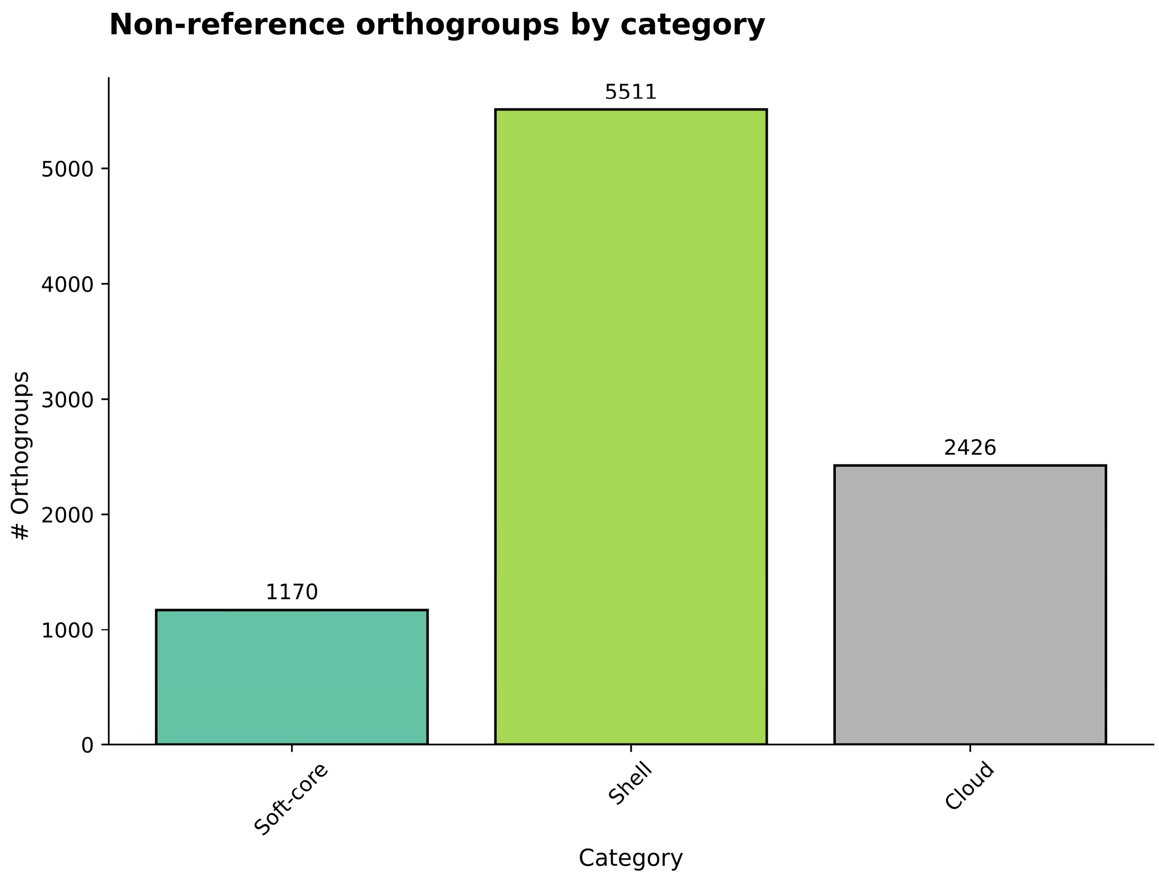


**Supplementary Figure 9. Non-reference orthogroups contributed by the *P. americana* accessions.** Bar plot showing the number of orthogroups that are absent from the West-Indian pure reference but present in at least one other assembly. Orthogroups are binned by the same pangenome frequency classes used elsewhere: Soft-core (present in 85 – 94% of haplotypes; turquoise), Shell (20 –84%; green), and Cloud (≤ 19%; grey); by definition, none of these "non-reference" orthogroups are Core. The y-axis reports the number of orthogroups in each class, with totals printed above bars. Orthogroups were inferred with OrthoFinder v2.5.5[8] (within GENESPACE v1.3.1[9]) and presence/absence summarised with pangenome_pav_support.py.

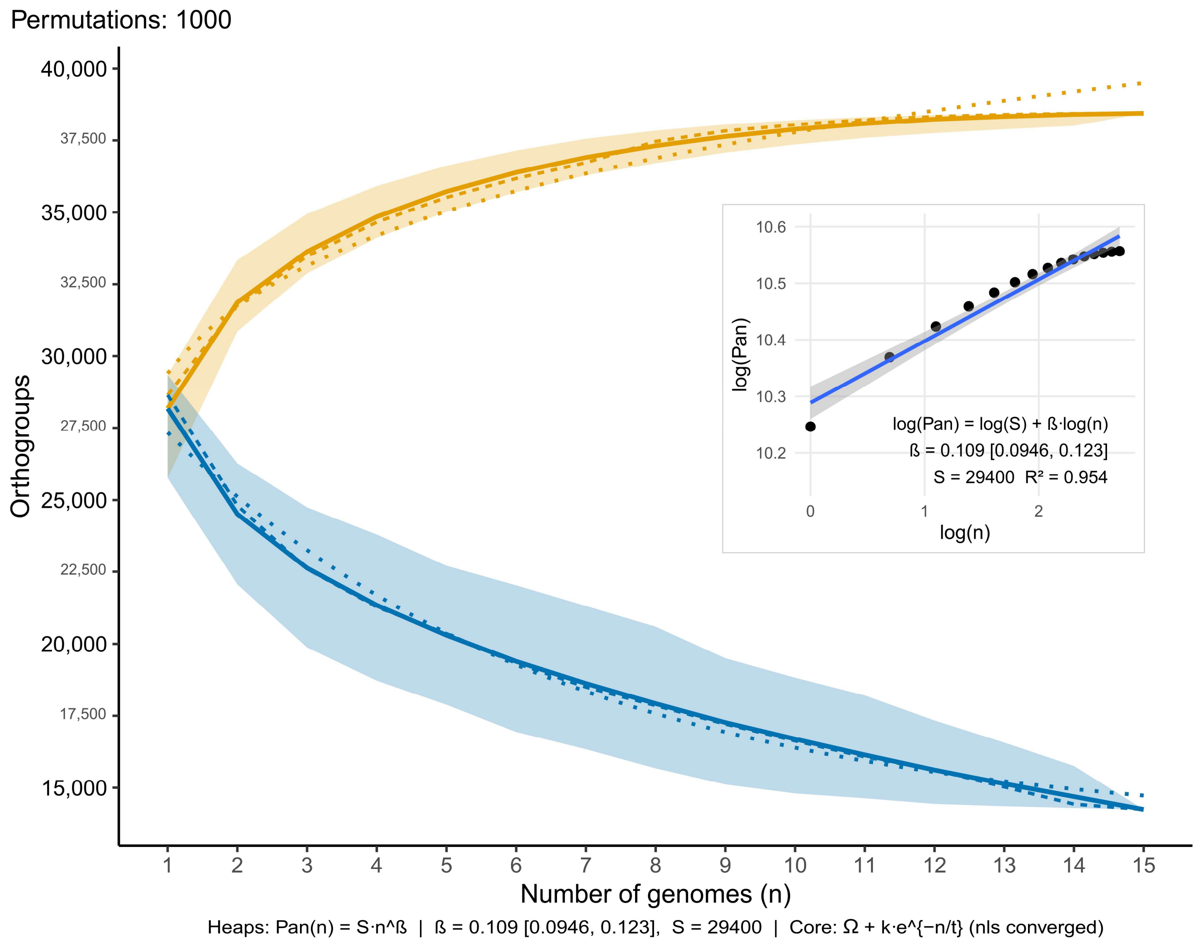


**Supplementary Figure 10. Pan- and core-genome orthogroup accumulation curves for 15 *Persea americana* haplotypes.** Curves were computed across 1,000 random permutations of genome order; bold lines show the mean number of orthogroups, dashed lines the median, and shaded ribbons the 95% confidence intervals. The dotted orange line is the Heaps' law fit for the pangenome, $Pan(n) = S{\cdot}n\text{\^{}}\beta$ (*n*, number of genomes; *S*, scale constant; $\beta$, openness exponent). The dotted blue line is the asymptotic model for the core-genome, $Core(n) = \Omega + k{\cdot}e\text{\^{}}(-n/\tau)$ ($\Omega$, asymptotic core size; *k*, amplitude; $\tau$, decay constant). Inset: log–log regression used to estimate $\beta$ ($\beta = 0.109$, 95% CI 0.0946 – 0.123; $S = 29{,}400$; $R^2 = 0.954$).

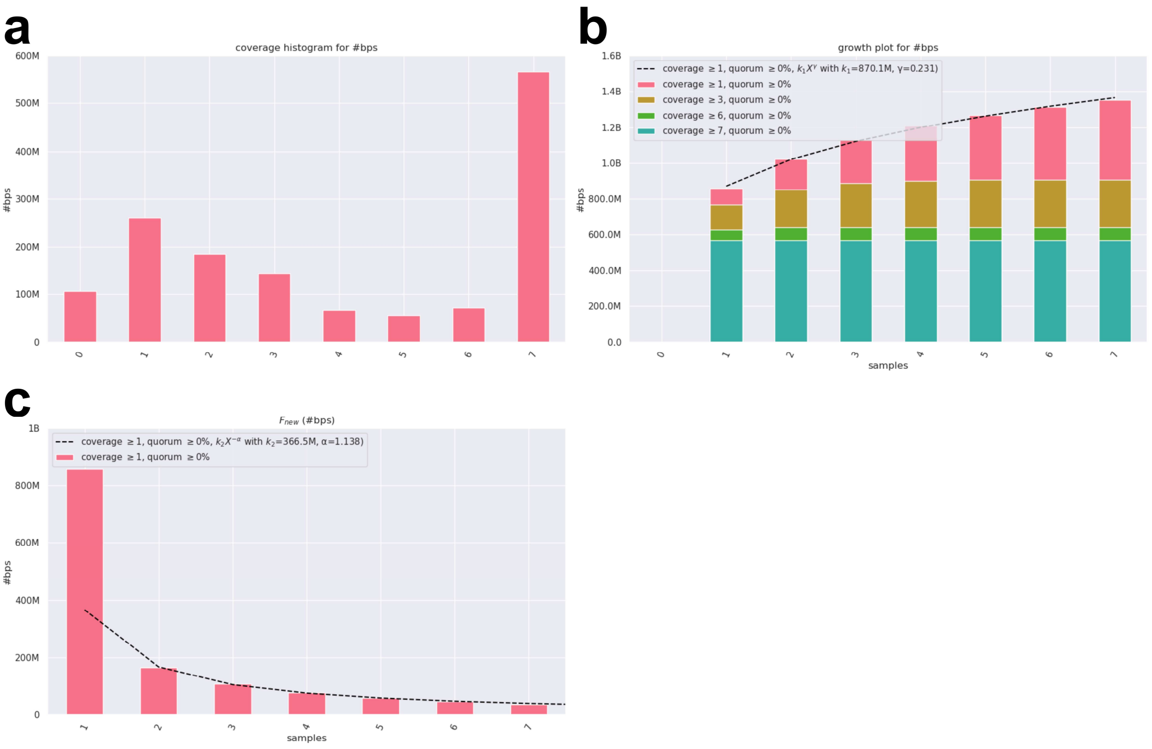


**Supplementary Figure 11. Sequence-level pan-genome growth from Panacus.** Panacus v0.3.5[10] was run on GFA walk records at the bp level (quorum ≥ 0% and coverage classes ≥1, ≥3, ≥6, ≥7, with sample merging for haplotype assemblies). **a**) Coverage histogram partitioning total sequence by coverage class (bases present in at least *k* assemblies). **b**) Cumulative pan-sequence growth as assemblies are added; stacked bars show the contribution of each coverage class, and the dashed line is the Heaps' law fit $k_1X^\gamma$ ($\gamma \approx 0.231$). **c**) New sequence contributed by each additional assembly; dashed line is the decay fit $k_2X^{-\alpha}$ ($\alpha \approx 1.138$), indicating that novel sequence discovery is slowing as the sequence space approaches saturation. Curves rendered with panacus-visualize.

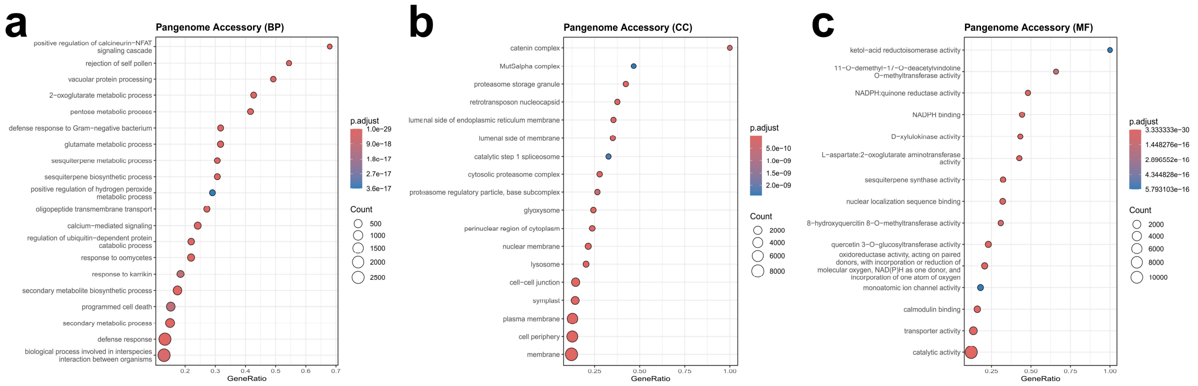


**Supplementary Figure 12. Gene Ontology enrichment of the pangenome accessory set.** Bubble plots summarizing GO terms enriched among accessory genes (Shell + Cloud orthogroups) across all assemblies. Enrichment was computed with topGO v2.58.0[11] (classic Fisher's exact test) separately for the **a**) Biological Process (BP); b) Cellular Component (CC); and c) Molecular Function (MF) ontologies, using all annotated genes as the background and restricting terms to the *Viridiplantae* lineage (GO taxon-constraints). *P*-values were corrected with Benjamini–Hochberg FDR; circles are coloured by adjusted *p*-value and scaled by the number of genes annotated to each term, while the x-axis shows GeneRatio (term count / accessory total). Within each ontology, redundant terms were reduced by semantic similarity (similarity cutoff = 0.7; clusterProfiler v4.14.6[12]) and results were visualized with enrichplot.

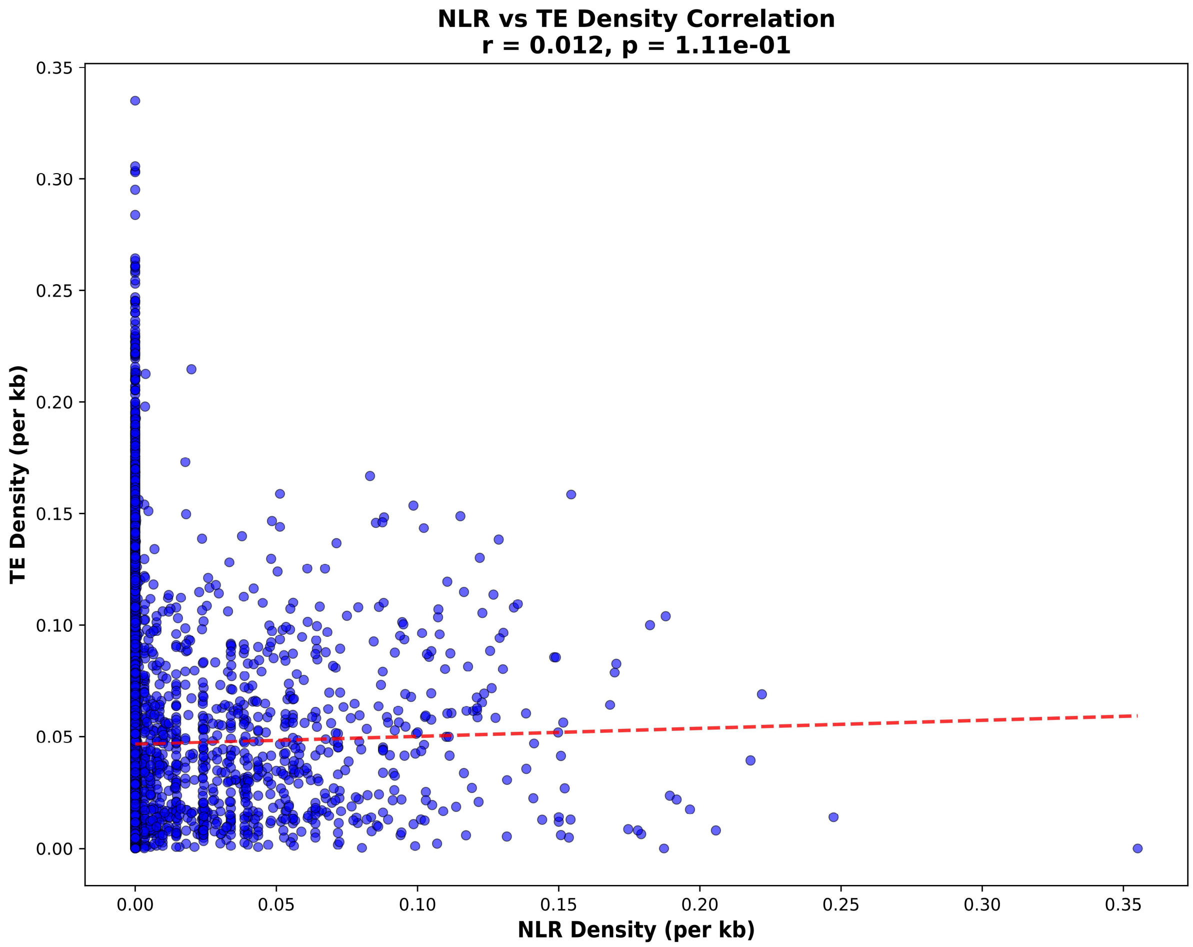


**Supplementary Figure 13. Correlation between Nucleotide-binding Leucine-rich repeat (*NLR*) genes and TE density across avocado genomes.** Scatter plot of *NLR* density (per kb) versus transposable element (TE) density (per kb), calculated from GFF3 annotations in 50 kb non-overlapping windows. Each point represents a genomic window. A linear regression trendline (red dashed) is shown, with the Pearson correlation coefficient (r) and associated *p*-value indicated. *NLR* and TE densities were calculated using a custom Python script (gff3_density_analyzer.py), which applies Gaussian smoothing to window-based counts.

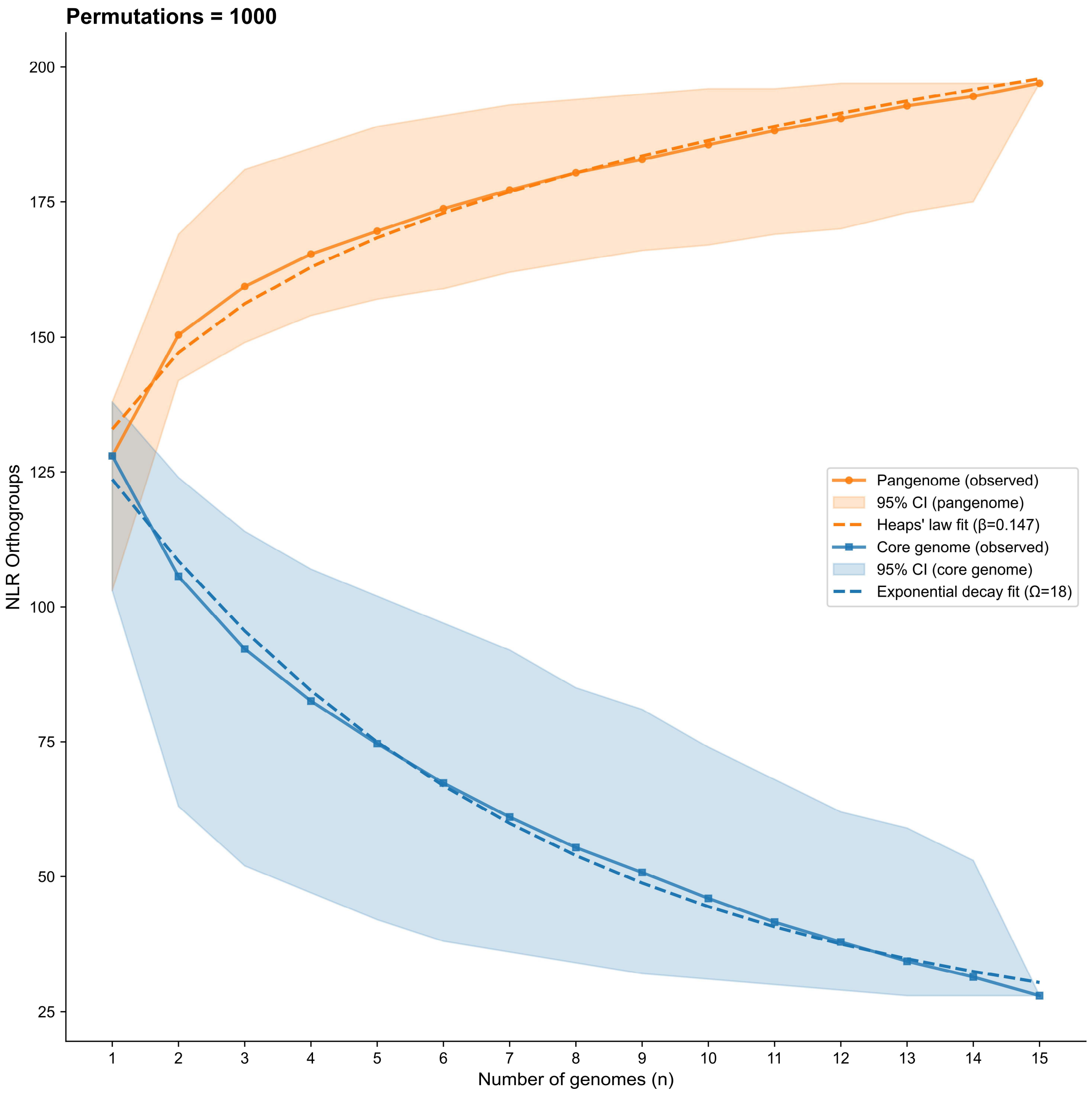


**Supplementary Figure 14. Pan- and core-Nucleotide-binding Leucine-rich repeat (*NLR*) orthogroup accumulation curves for 15 avocado haplotypes.** Accumulation curves were computed from GENESPACE orthogroup assignments across 1,000 random permutations of accession order. Bold lines show the mean number of orthogroups, dashed lines the median, and shaded ribbons the 95% confidence intervals. The dotted orange line is the Heaps' law fit for the NLR pangenome, $Pan(n) = S \cdot n^\beta$ (n, number of accessions; S, scale constant; β, openness exponent). The dotted blue line is the asymptotic model for the NLR core, $Core(n) = \Omega + k \cdot e^{(-n/\tau)}$ (Ω, asymptotic core size; k, amplitude; τ, decay constant).
